# A modality independent proto-organization of human multisensory areas

**DOI:** 10.1101/2022.03.14.484231

**Authors:** Francesca Setti, Giacomo Handjaras, Davide Bottari, Andrea Leo, Matteo Diano, Valentina Bruno, Carla Tinti, Luca Cecchetti, Francesca Garbarini, Pietro Pietrini, Emiliano Ricciardi

**Affiliations:** MoMiLab, IMT School for Advanced Studies Lucca, Lucca, Italy; Department of Translational Research and Advanced Technologies in Medicine and Surgery, University of Pisa, Pisa, Italy; Department of Psychology, University of Turin, Turin, Italy

## Abstract

The processing of multisensory information is based upon the capacity of brain regions, such as the superior temporal cortex, to combine information across modalities. However, it is still unclear whether the representation of coherent auditory and visual events does require any prior audiovisual experience to develop and function. In three fMRI experiments, intersubject correlation analysis measured brain synchronization during the presentation of an audiovisual, audio-only or video-only versions of the same narrative in distinct groups of sensory-deprived (congenitally blind and deaf) and typically-developed individuals. The superior temporal cortex synchronized across auditory and visual conditions, even in sensory-deprived individuals who lack any audiovisual experience. This synchronization was primarily mediated by low-level perceptual features and relied on a similar modality-independent topographical organization of temporal dynamics. The human superior temporal cortex is naturally endowed with a functional scaffolding to yield a common representation across multisensory events.

## INTRODUCTION

The ability to combine signals across different sensory modalities is essential for an efficient interaction with the external world. To this end, the brain must detect the information conveyed by different sensory inputs and couple coherent events in space and time (i.e., solve the correspondence problem^1^). Specifically, when processing audiovisual information, signals from sight and hearing converge onto multiple brain structures and, among them, the superior temporal cortex is acknowledged as being a pivotal hub^2,3^. Evidence exists that basic multisensory processing is already present in newborns^4^, while audiovisual experience appears to be critical for the development of more complex multisensory computations lifelong^5,6^. Nonetheless, the extent to which audiovisual experience is a mandatory prerequisite for the superior temporal cortex to develop and become able to detect shared features between the two sensory streams is still undefined. Adult individuals who specifically lack visual or auditory input since birth represent an optimal model to test whether brain computations do require a complete audiovisual experience to develop^7,8^.

Here we determined the synchronization of brain responses in two groups of sensory-deprived adults (SD - i.e., congenitally blind and deaf) and in three samples of typically-developed (TD) individuals exposed to the audiovisual, audio-only or visual-only versions of the same long-lasting narrative. This approach, called Intersubject Correlation (ISC) analysis, postulates that brain regions do synchronize across individuals when processing the same stimulus features^9^. Therefore, any evidence of synchronization within the superior temporal cortex both across conditions and experimental groups, would be indicative that this region yields shared representations of visual and auditory features despite so different postnatal sensory experiences. Furthermore, we provided a thorough description of the events occurring across the visual and auditory streams by developing a model-mediated version of ISC. This approach determined whether brain synchronization resulted from the processing of coherent low-level visual (e.g., motion energy) and acoustic (e.g., spectral properties) features, or it was instead driven by high-level semantic (e.g., language and story synopsis) characteristics. Finally, additional analyses characterized the temporal dynamics of the synchronization across individuals and depict the chronotopic organization of multisensory regions.

As expected, the activity of the superior temporal cortex was synchronized across auditory and visual inputs in TD participants. Crucially, the synchronization was also present across SD individuals, despite the congenital lack of any auditory or visual input since birth and the distinct postnatal experiences. Furthermore, the synchronization was mediated by low-level perceptual features in both TD and SD groups and relied on a similar modality-independent topographical organization of temporal dynamics consisting of adjacent cortical patches tuned to specific receptive windows. Altogether, these observations favor the hypothesis that the human superior temporal cortex is naturally endowed with a functional scaffolding to yield a common neural representation across coherent auditory and visual inputs.

## RESULTS

The ISC analysis^9^ was used to measure the similarity in the brain responses elicited by the processing of either the audiovisual, the auditory or the visual streams of the same naturalistic narrative (i.e., the live-action movie *101 Dalmatians*) presented to both TD and SD participants by means of three fMRI experiments. In addition to the overall measure of synchronization provided by the ISC approach, we built a hierarchical set of models describing the low-level and the high-level features of the movie auditory and visual streams to test which stimulus properties mediate the interaction across senses. Finally, the temporal properties (i.e., the temporal receptive window^10^) of the dynamical processes responsible for the synchronization across senses were studied and compared in the three experiments.

In a first experiment, the neural correlates of the audiovisual (AV) stimulus were studied in a sample of TD participants to establish how the brain processes multisensory information. In a second experiment, two unimodal versions of the same movie (i.e., visual-only - V -, and auditory-only - A -) were created by excluding one or the other sensory channels. Then, to investigate to what extent the neural representation of the same narrative is shared across sensory modalities, the similarity between visually- and auditory-evoked brain responses (*A vs V*) was evaluated by performing ISC analysis across the two samples of TD participants who were exposed to either A or V conditions. Finally, in a third experiment, we studied the brain synchronization between blind and deaf participants (*A vs V*), listening to the audio-only (A) and watching the visual-only (V) movie, respectively.

### Synchronized brain responses between vision and hearing in typical developed individuals

In the first experiment (Figure 1A), a whole-brain ISC analysis was computed on TD participants exposed to the audiovisual version of the narrative (i.e., multisensory condition, within-condition ISC, n=10). The statistical significance of synchronization maps was based on non-parametric permutation tests and Family-Wise-Error (FWE) correction was applied (p<0.05). As shown in Figure 2A, results highlighted a set of regions involved in the processing of multisensory information, encompassing a large extent of the cortex (~40% of the cortical volume). Significant synchronized regions included primary sensory regions, such as early auditory and visual areas, as well as high-order cortical areas, such as the Superior Temporal Gyrus (STG) and Sulcus (STS), the inferior parietal region, the precuneus, the posterior and anterior cingulate cortex (PostCing and AntCing, respectively), the inferior frontal gyrus, as well as the dorsolateral and dorsomedial portions of the prefrontal cortex. The ISC peak was found in the central portion of the left STG (peak r=0.452, 95^th^ range: 0.206:0.655; MNI: −65 −14 +1).

**Figure 1.**
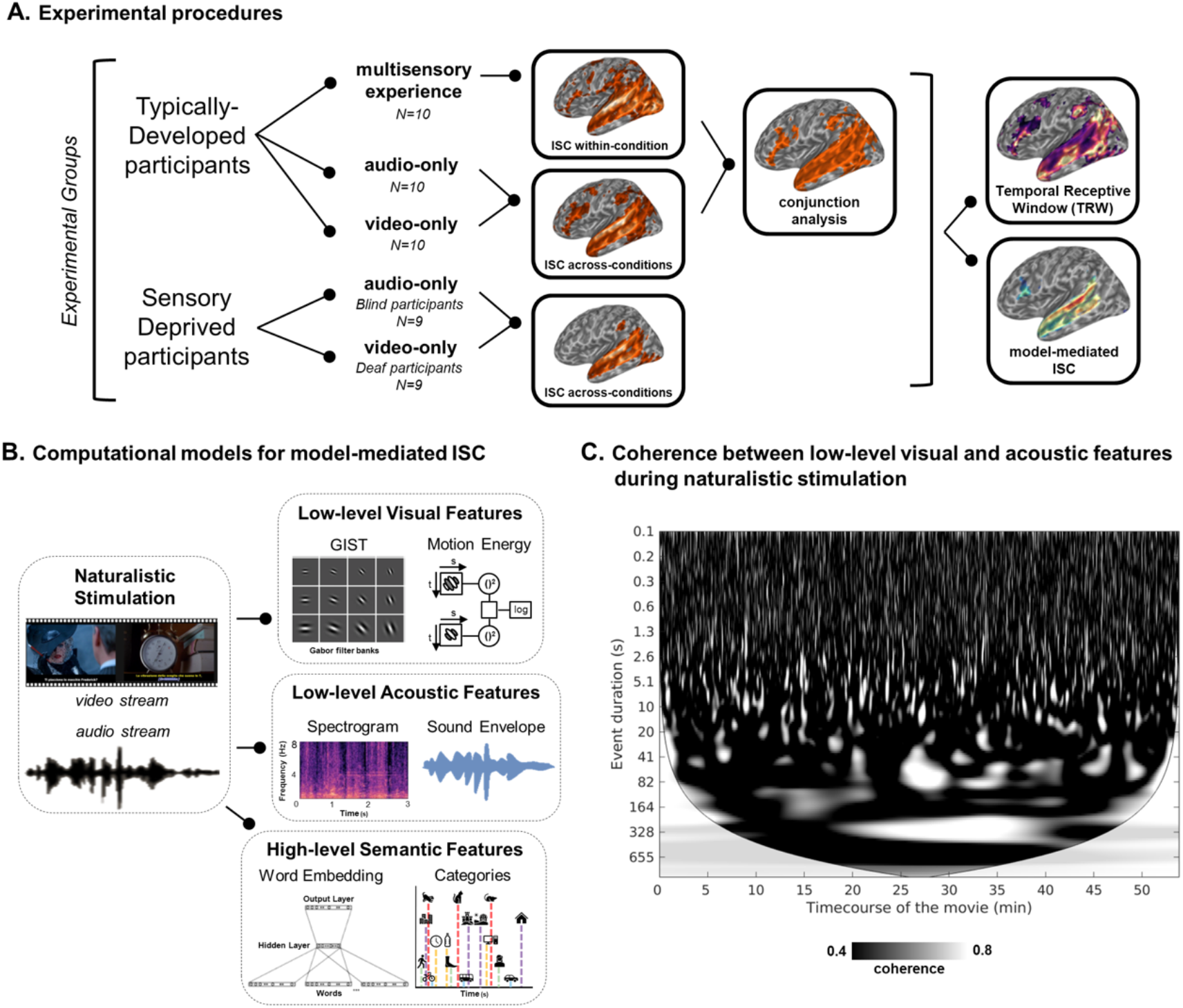
In Panel A, all the experimental conditions, procedures and analyses are reported. In a first experiment, the neural correlates of a full audiovisual (AV) stimulus were studied in a sample of TD participants to examine how the brain processes these types of multisensory information. In a second experiment, two unimodal versions of the same movie (i.e., visual-only -V-, and auditory-only -A-) were presented and the similarity across visually- and auditory-evoked brain responses (*A vs V*) was assessed in two samples of TD participants. In a third experiment, the role of audiovisual experience for the emergence of these shared neural representations was tested in congenitally SD individuals by measuring the similarity of brain responses elicited across blind and deaf participants. In Panel B, we provide a brief description of the features extracted through computational modeling from the movie. Movie-related features fall into two categories: i) low-level acoustic and visual features; and ii) high-level semantic descriptors. The first category comprises fine-grained features extracted from the auditory (e.g., spectral and sound envelope properties to account for frequency- and amplitude-based modulations) and visual streams (e.g., set of static Gabor-like filters and motion energy information based on their spatiotemporal integration). These features showed a variety of temporal dynamics and were of utmost interest for our analysis since they encoded collinearities across the two streams during naturalistic stimulation. As concerns the set of high-level features, we performed manual annotation of both visual (e.g., a close-up of a dog) and sound-based (e.g., barking of a dog) natural and artificial categories (see Supplementary Methods for more details), and we exploited machine learning techniques (e.g., word embedding) to define a group of descriptors representing the semantic properties of the auditory and visual streams (see Supplementary Materials for further details). In Panel C, we show the results of a continuous wavelet transform analysis applied to the movie acoustic and visual signals to evaluate the existence of collinearities across the low-level features of the two sensory streams. Figure depicts the presence of cross-modal correspondences, with hundreds of highly coherent events (white marks) distributed along the time course of the movie (x axis), lasting from a few tenths of a second to several minutes (y axis).

**Figure 2.**
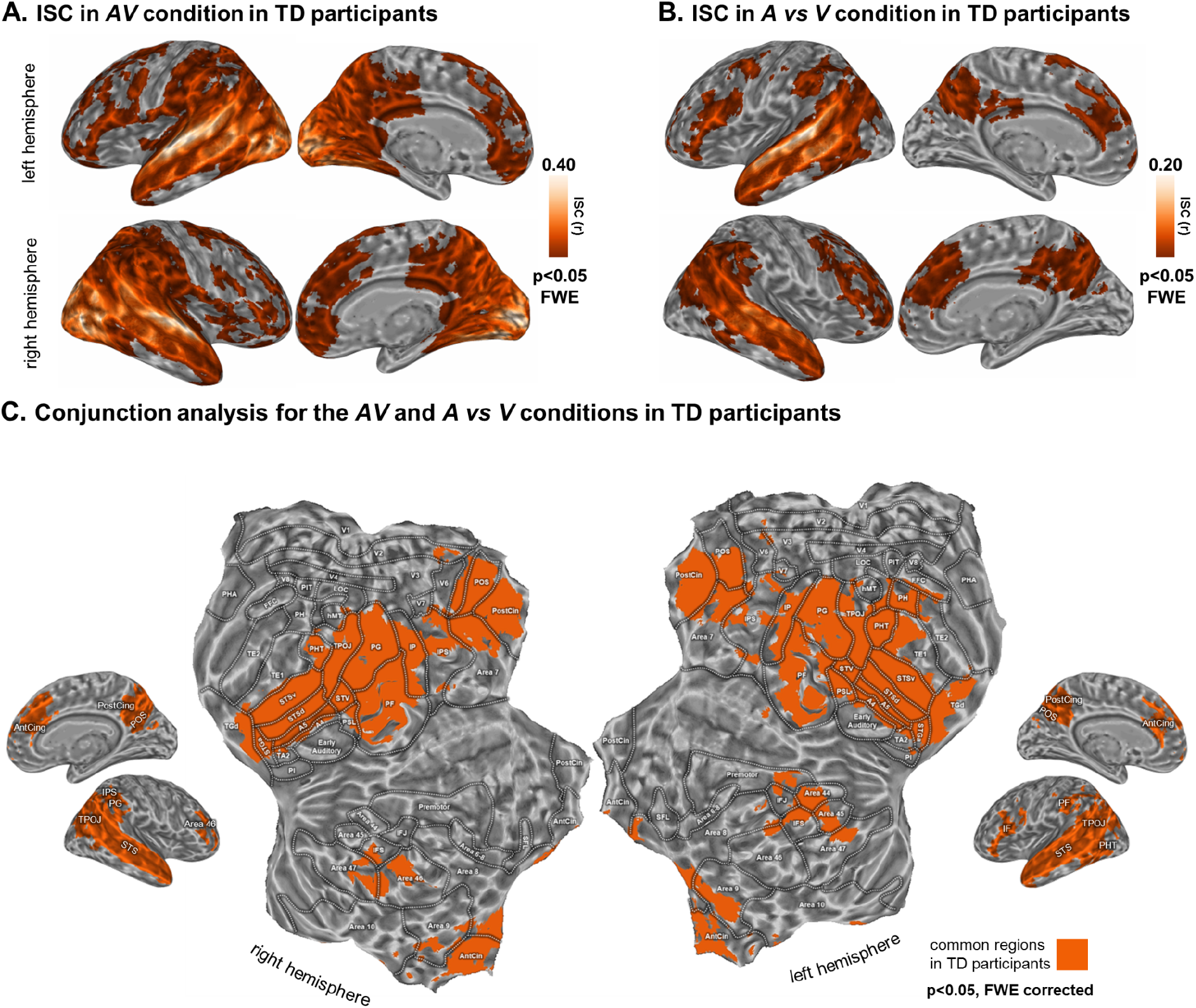
In Panels A and B, we report the ISC in TD participants in the *AV* and *A vs V* conditions respectively. In Panel C, the conjunction analysis of the two aforementioned experimental conditions is shown.

The second experiment measured the interplay between vision and audition in two groups of TD individuals (n=10 A-only, n=10 V-only, Figure 1A). Synchronization (*A vs V*, across-modalities ISC, Figure 2B) was present in the ~14% of the cortical volume with no involvement of primary auditory and visual areas. Significant regions comprised the superior portion of the temporal lobe, inferior parietal, precuneus, cingulate and prefrontal cortical areas. As in the case of the AV modality, the synchronization across A-only and V-only conditions peaked in the left central portion of STS (peak r=0.214, 95^th^ range: 0.054:0.485; MNI: −65 −32 +1).

The brain areas identified from the above experiments in TD participants were targeted in a conjunction analysis (p<0.05, FWE corrected, Figure 2C) to highlight regions synchronized during the multisensory experience, which also shared a common neural representation across the A-only and V-only conditions. To provide a finer anatomo-functional characterization of brain regions included in the conjunction map, we adopted the Human Connectome Project (HCP) parcellation atlas^11^ (please refer to the Supplementary Table 1 for a detailed description of cortical labels). The map (Figure 2C, 3B) identified a set of six cortical regions, which were commonly recruited across the A-only, V-only and multisensory experimental conditions in TD participants. Specifically, the highest degree of spatial overlap was found in a bilateral temporo-parietal cluster, which comprised the superior temporal cortex (A4, A5, STSd, STSv, STGa, TGd, STV, PSL), portions of the ascending branch of the inferior temporal gyrus (PHT), the temporo-parieto-occipital junction (TPOJ) and the inferior parietal cortex (PG, PF, IP). Two additional bilateral clusters were identified: the first located in the posterior parietal cortex, comprising the PostCing, the parieto-occipital sulcus (POS), the superior parietal (corresponding to Brodmann Area-BA 7), and the second in the medial prefrontal cortex, enclosing the bilateral AntCing and the dorsomedial portions of the superior frontal gyrus (BA 9). Lastly, two lateralized clusters were found in the left inferior frontal gyrus (BA 44 and 45) and in the right prefrontal cortex (BA 46 and 47). Of note, the conjunction map did not reveal any early sensory areas indicating that the activity of those regions was not synchronized in response to the audio-only and visual-only conditions (*A vs V*, Figure 2B).

**Figure 3.**
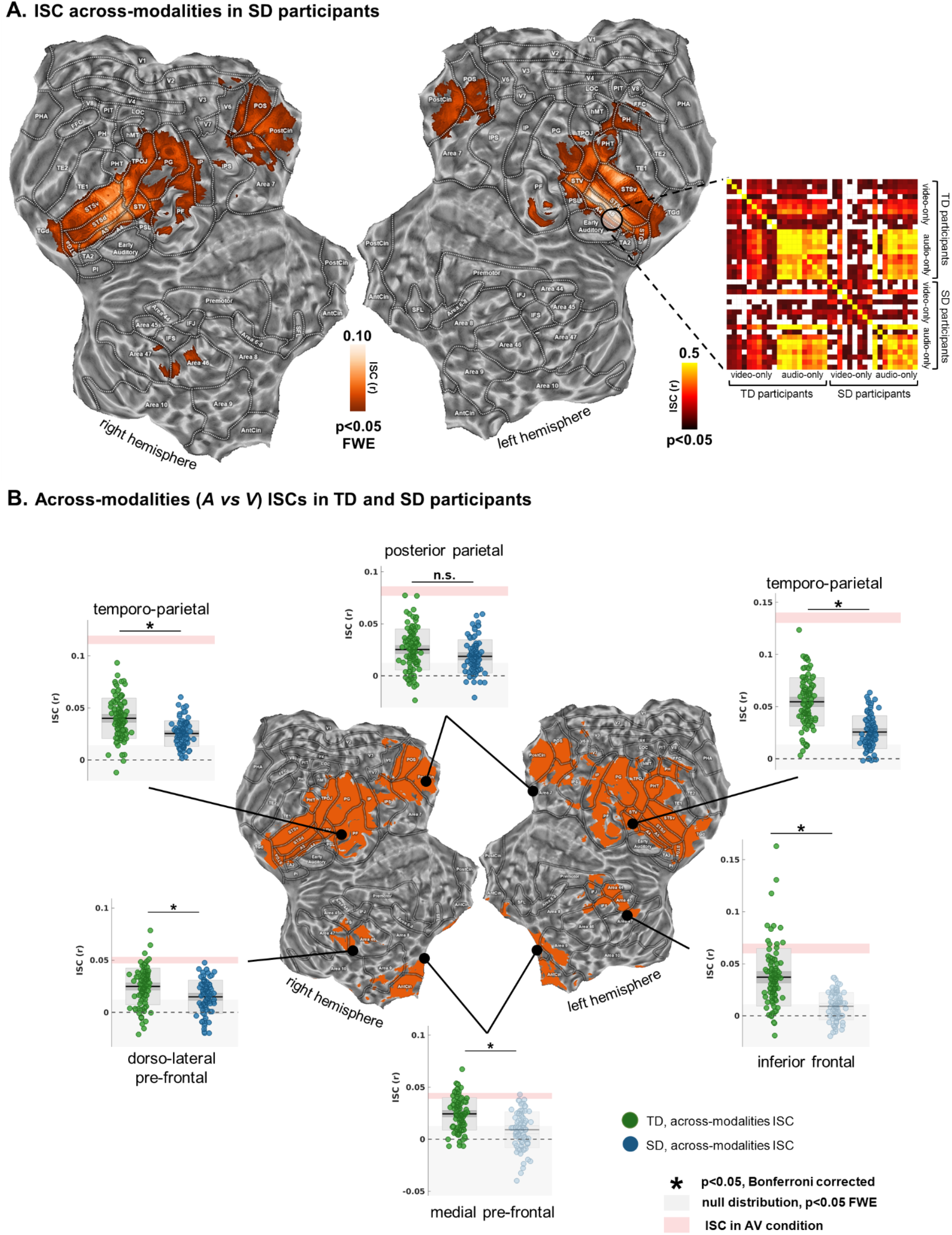
Panel A shows the results from the across-modality ISC in SD participants (p<0.05, FWE corrected). In the matrix on the right, the raw ISC across TD and SD participants was reported for the A-only and V-only conditions. ISC magnitude (Pearson’s r coefficient) was extracted from a region of interest (6 mm radius) centered in the left STG, around the peak of ISC of the first experiment. White cells indicated subject pairings below the significant threshold (p<0.05, uncorrected for multiple comparisons). In Panel B, figure reports the ISC magnitude in the three experimental groups: for each brain region obtained from the conjunction analysis (six clusters, minimum cluster size of 20 adjacent voxels), the *A vs V* condition in TD participants was compared to *A vs V* in SD individuals. Moreover, the average ISC (with its standard error) in the *AV* condition is reported, with a shaded area in rose, as a ceiling effect due to multisensory integration. Transparency is applied to the color of points and boxplots to indicate that the group ISC was not significant when compared to a null distribution. In each boxplot, the dark horizontal line represents the sample mean, the dark-grey shaded box the 95% confidence intervals of the standard error of the mean, and the lightgrey shaded box reports the standard deviation. Multiple comparisons were handled using Bonferroni correction (p<0.05).

Altogether, results of experiments 1 and 2 showed that well-known multisensory areas^2^ synchronize to audiovisual correspondences over time, even when sensory features are provided unimodally.

### Synchronized brain responses between vision and hearing in congenitally deprived individuals

In the third experiment, ISC analysis tested the similarity of brain responses across V-only and A-only conditions in congenitally deaf and blind participants respectively. Specifically, across-modality ISC (i.e., *A vs V;* Figure 1A) was performed in nine blind and nine deaf individuals. In this experiment, ISC was computed in the regions of interest identified by the conjunction map obtained from the first two experiments with TD participants.

As shown in Figure 3A, congenital lack of either auditory or visual experience did not prevent synchronization of brain responses across modalities. Indeed, significant synchronization was found in the bilateral temporal cortex (A4, A5, STS, STV, STGa), TPOJ, PostCing and POS, which represented ~47% of the conjunction map identified in TD participants and 5% of the overall cortical volume (p<0.05, FWE corrected). Moreover, the ISC map highlighted a minimal involvement of the bilateral inferior parietal (PG and PF), the right dorsolateral prefrontal (BA 46) (~1% of the conjunction mask), and the left inferior temporal cortex (PHT and PH; ~1%). Notably, SD individuals showed the ISC peak within the central portion of left STS similarly to TD participants (peak r=0.131, 95^th^ range: −0.027:0.372; MNI: −62 −32 +1). On the contrary, SD groups did not show any significant synchronization in bilateral medial prefrontal (AntCing, BA 9), left inferior frontal (BA 44 and 45), and prefrontal cortex (Area 46 and 47). To further investigate the consistency of synchronization at single subject level, the raw ISC was computed across all participants in a region of interest defined in the left STG (Figure 3A). The ISC matrix confirmed high synchronization between subject pairings of the audio-only conditions, in line with the role of the temporal cortex in auditory computations. Additionally, spread synchronizations emerged between the individuals exposed to the visual-only condition and all the other participants, supporting the hypothesis of a modality-independent processing of information in this cortical patch.

The comparison between TD and SD participants in the *A vs V* condition (Figure 3B) showed a diffuse decrease of ISC across all the explored regions (Wilcoxon rank sum test, p<0.05, Bonferroni corrected), with the notable exception of the posterior parietal cortex (p>0.05). Moreover, in SD participants, the left inferior frontal gyrus and the bilateral medial prefrontal cortex were not statistically synchronized, with averaged ISCs falling within the null distribution.

Altogether, these results indicated that congruent auditory and visual streams elicited a functional synchronization in the superior temporal areas and in the postero-medial parietal cortex even in the case of congenital auditory or visual deprivation and thus, in absence of prior audiovisual experience.

### Do synchronized brain responses rely on perceptual or semantic stimulus features?

Brain activity of congenitally deaf and blind people was synchronized when exposed to the same narrative. Nevertheless, whether the synchronization could be ascribed either to the processing of perceptual (i.e., low-level) features, or to semantic (i.e., high-level) representations shared across the different conditions, remained to be determined. We took advantage of computational modeling (Figure 1B) to extract fine-grained, low-level features from both the auditory (e.g., spectral and sound envelope properties to account for frequency- and amplitude-based modulations) and visual streams (e.g., set of static Gabor-like filters and motion energy information based on their spatiotemporal integration). Moreover, a set of high-level features was collected by means of manual annotation and automated machine learning techniques (e.g., word embedding) to represent the linguistic and semantic properties of the narrative.

To address the role of low-level visual-acoustic and high-level semantic features, we measured the impact of each model in modulating the magnitude of the ISC between the unimodal conditions (Figure 4; p<0.05, FWE corrected). Specifically, each model was regressed out from the individual brain activity before computing the ISC. This procedure resulted in a reduction of the ISC value that reflects, for each brain region, the relevance of distinct stimulus descriptors that were regressed out. Therefore, the role of each model as a *mediator* was evaluated on the synchronization of brain responses across participants and the relative drop in the ISC magnitude was computed for each voxel^12^. In principle, if a model explains entirely the activity of a specific brain region, the ISC in that area will drop substantially, with values that will approach zero. Conversely, if a model does not contribute to the synchronization of brain responses across visual and auditory movie processing, the drop in the ISC magnitude will be negligible. We named this approach *‘model-mediated ISC’*.

**Figure 4.**
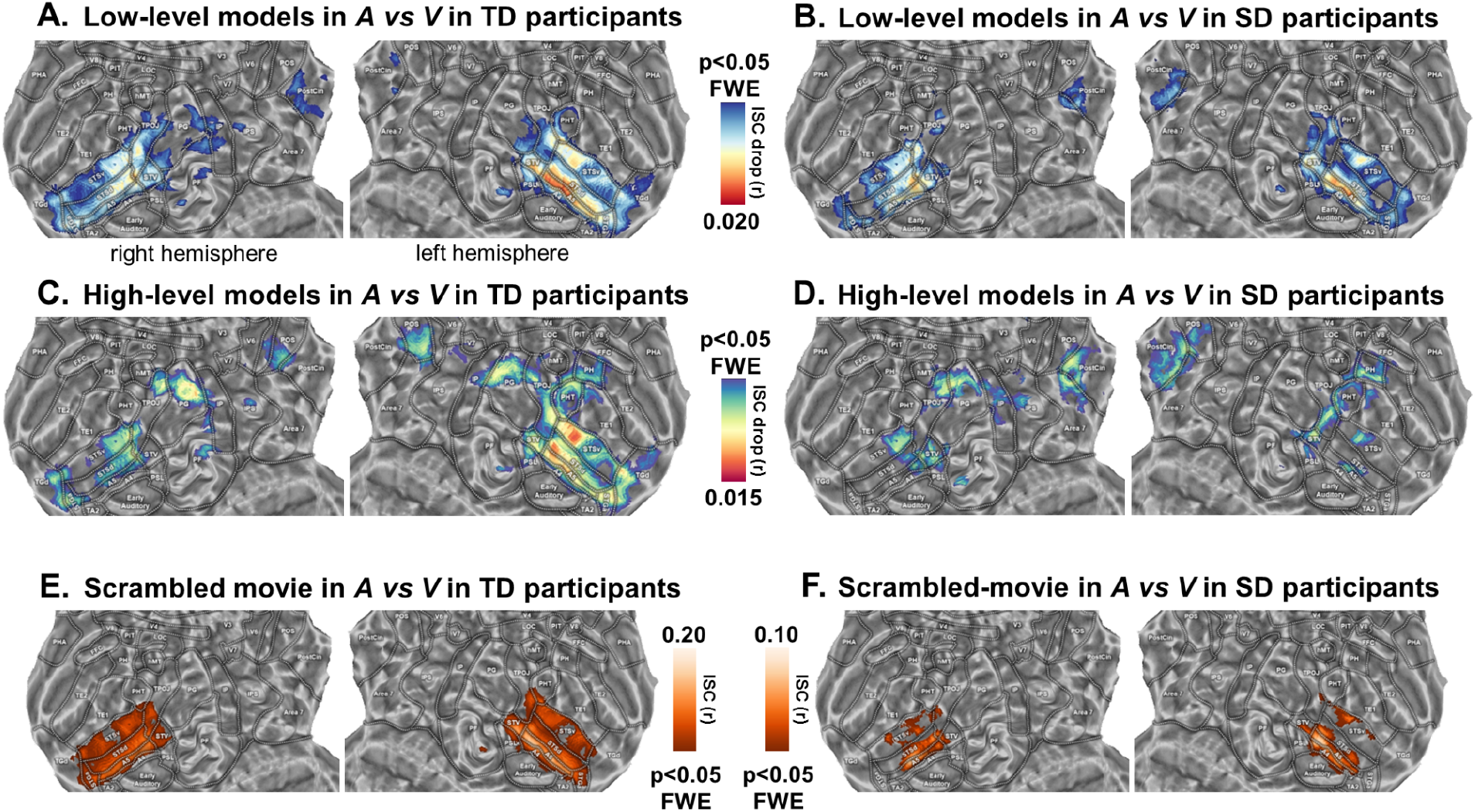
In Panels A and B, we report the model-mediated ISC across TD and SD in the *A vs V* condition for the low-level models, based on the movie acoustic and visual properties. In Panels C and D, we depict the model-mediated ISC across TD and SD individuals in the *A vs V* condition for the high-level models, based on semantic features. In Panels E and F, we show the ISC across TD and SD individuals in the *A vs V* condition during the processing of the scrambled movie. Note that, only the temporo-parietal cortex is mapped, since the results in frontal areas were present just for the TD participants. All results were corrected for multiple comparisons (p<0.05, FWE).

Concerning the low-level models, we regressed out visual features from brain activity during the A-only stimulation and acoustic features from the activity during the V-only processing to test whether a drop of ISC magnitude could be ascribed to audiovisual correspondences. Thus, this procedure identifies the impact of the unique portion of model variance shared across the two modalities. In both TD and SD groups the drop of ISC was significant in the posterior parietal and STS/STG regions (A4, A5, STSd), and maximum in the central portion of the left STS (Figure 4A-B; drop of ISC at peak: TD *A vs V:* r=0.020, 95^th^ range: 0.006:0.039, MNI: −62 −26 1; deprived *A vs V:* r=0.018, 95^th^ range: −0.014:0.061, MNI: −65 −26 1). Consequently, these cortical areas retain a low-level representation of audiovisual features that inherently co-occur in a naturalistic stimulation.

Finally, the role of language and semantics was investigated combining the features generated by the representation of the stream of words and those extracted by manual annotation of categories. Therefore, these high-level features, which are naturally multimodal, were removed from the brain activity of all participants. Results of the model-mediated ISC revealed that semantic features had a significant impact in synchronization across modalities in the posteromedial parietal cortex in both TD and SD participants. As regards the temporal cortex, in SD participants only the superior temporal visual area (STV) and the posterior portions of STS were affected by regressing out semantic features from brain activity, whereas in TD participants model influence was spread across the whole STS, and particularly in the left hemisphere (Figure 4C-D; drop of ISC at peak: TD *A vs V* in left mid-STS: r=0.015, 95^th^ range: −0.001:0.036, MNI: −53 −35 1; deprived *A vs V* in the right TPOJ: r=0.009, 95^th^ range: −0.010:0.034, MNI: 49 −74 19).

These results demonstrate that, in both TD and SD, STG/STS synchronization was primarily driven by lower-level properties, whilst high-level semantic features are represented within the same brain region in TD individuals only.

### Do synchronized brain responses depend on changes in the movie plot?

Model-mediated analyses clarified the contribution of low- and high-level features to brain synchronization. However, this approach did not test the extent to which the temporal sequence of connected events in the narrative (i.e., story synopsis) determines brain synchronization. To account for this possible mechanism, a control condition was based on a scrambled short movie in which we manipulated the chronological order of previously unseen cuts (lasting from 1 to 15 seconds, median: 3.5 seconds, maintaining the same distribution of cut lengths of the original movie). Because of this manipulation, even though the storyline of the control condition was nonsensical, a set of stimulus features were preserved: (1) the cinematography, with the same coarse- and fine-grained visual features, (2) the sound mixing, (3) a semantic representation based on single words up to very short sentences, and (4) the editing pacing. Importantly, we left untouched the congruency between audio and visual streams, which remained synchronized with each other.

Interestingly, the scrambling of the movie plot differentially affected the synchronization across brain areas. Indeed, a meaningless narrative still triggered shared responses in the central and posterior parts of superior temporal cortex (*A vs V* in TD and SD), particularly in A4, A5 and STSd, and the peak was located in the central part of the left STS, with similar intensities as compared to the original movie (ISC at peak: TD *A vs V:* r=0.162, 95^th^ range: −0.004:0.383, MNI: −64 −14 −4; SD *A vs V:* r=0.105, 95^th^ range: −0.050:0.397, MNI: −64 −32 2). Remarkably, the disruption of the narrative significantly affected the posteromedial parietal regions (Figure 4E-F), whose synchronicity did not reach the significance threshold in both TD and SD individuals.

Therefore, this further evidence confirms that synchronous correlations in specific portions of the temporal cortex (A4, A5, STSd) were primarily driven by low-level perceptual properties and not by high-level semantic computations required for the processing and understanding of the narrative.

### Temporal dynamics of the synchronized brain responses between vision and hearing

Additional analyses were conducted to characterize the temporal dynamics of the synchronization across individuals. Our aim was to test whether each experimental condition retained the same temporal dynamics and to describe the diachronic properties of the explored cortical regions.

First, we evaluated the correspondences between the auditory and visual streams in our naturalistic stimulation. To compare visual (i.e., pixel intensities) and acoustic (i.e., sound wave energy) information, a set of descriptors (i.e., static Gabor-like filters and spectral features) were extracted at the highest available sampling frequency (25Hz, the frame rate of the original movie). Afterward, the coherence in time of the two streams was measured by means of a continuous wavelet transform to detect both the duration and the onset time of specific events in the movie, shared across auditory and visual streams. Although a set of relatively coarse computational features were used, the results reported in Figure 1C demonstrated the existence of a multifaceted series of highly coherent events, lasting from tenths of a second to several minutes. Considering the high variability in temporal dynamics of the correspondences across modalities, we expected that different brain regions would have expressed different temporal tunings to process the incoming sensory input. To address this point, we estimated the ISC in our regions of interest by means of a temporal filter of increasing window widths, from one time point (i.e., 2 seconds, which led to the same results of a classical ISC pipeline), up to four minutes. Being conceptually analogous to other techniques^10^, this methodological approach was defined as Temporal Receptive Window (TRW) analysis. We estimated both the length of the temporal window which retained the highest ISC (Figure 5, on the left) and the temporal profile of ISC for all the explored window widths (Figure 5, on the right). A high ISC over a short window would suggest that the correlation was modulated by rapidly changing events, whereas high ISC values over longer segments indicated that the correlation mostly relied on accumulating information.

**Figure 5.**
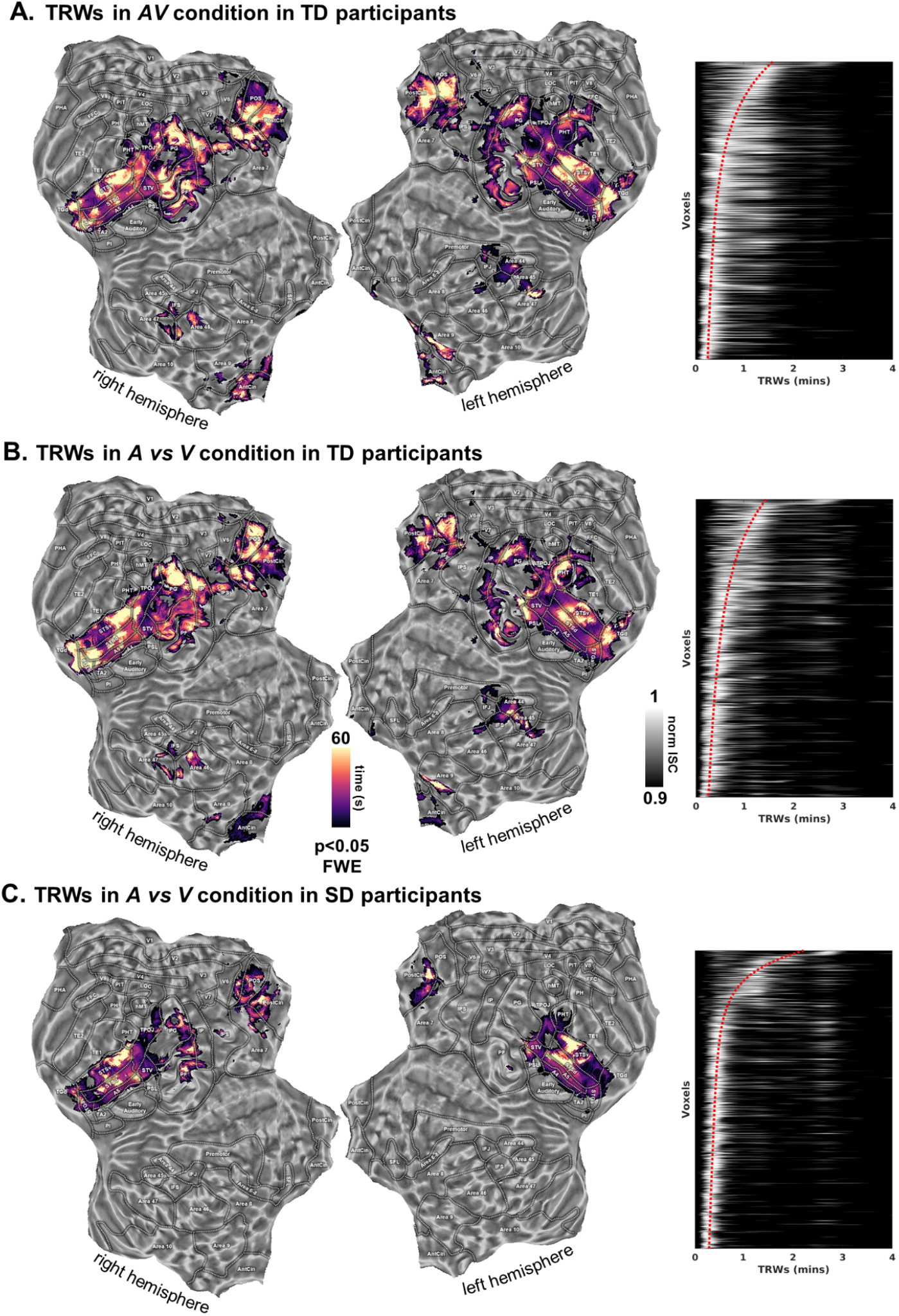
The figure shows a time-based representation of naturalistic information processing using sliding windows of different widths (from 2 secs to 240 secs) in the three groups (Panel A, B and C). On the left side of the figure, flat brain maps show the peak responses for the Temporal Receptive Windows (TRWs) in our regions of interest. Matrices on the right side indicate the overall synchronization profile across all the explored voxels which survived the statistical threshold (p<0.05, FWE corrected) in the *A vs V* condition in SD participants. The voxels were sorted according to the peak of the TRWs in the SD condition. Voxel order in the matrices was then kept constant across the three experiments. Pixel intensity depicts the normalized ISC (i.e., scaling the maximum to one) across all voxels and temporal windows. Red dotted lines represent the interpolated position of peaks across ordered voxels. Results showed that only a few voxels presented responses characterized by multiple peaks, whereas the majority demonstrated a clear preference for a specific temporal window and, more importantly, that the overall TRW profile was consistent across the three experimental procedures.

TRW results further highlighted the role of the superior temporal and posteromedial parietal cortex (p<0.05, FWE corrected) in detecting commonalities across multiple sensory streams. Specifically, coherent TRW maps across the three experimental conditions were found in the superior temporal cortex: A4, A5 and STS exhibited a patchy organization, in which adjacent subregions demonstrated distinct peak responses of temporal preference (Figure 5A, B and C, on the left). In details, ISC in A4, A5 and STSd showed a selective tuning for the fastest synchronization timescale, peaking at 10-20 seconds, whereas the middle and anterior portions of the sulcus, particularly in their ventral part, displayed a preference for timescales longer than one minute. Moreover, synchronization in the PostCing cortex and the POS was characterized mostly by slow modulations with an average preferred response to events occurring about every minute.

To summarize, the overall TRW profile was consistent across the three experimental procedures, indicating that synchronized regions retained a similar chronotopic organization.

## DISCUSSION

The present study questioned whether brain regions involved in audiovisual processing, including the superior temporal and neighboring areas, retain the ability to represent sensory correspondences across modalities, despite the complete lack of any prior auditory or visual experience. To this purpose, we characterized the coherence of neural responses evoked during a naturalistic stimulation across different modalities and across independent samples of typically developed, congenitally blind and congenitally deaf individuals, by performing three distinct fMRI experiments. Synchronized neural responses, selectivity for perceptual features and a preserved mapping of temporal receptive windows demonstrate that the functional architecture of the superior temporal cortex emerges despite the lack of audiovisual inputs since birth and irrespectively of postnatal sensory experiences. This observation favors the hypothesis that the human superior temporal cortex is endowed at birth with a functional scaffolding to yield basic audiovisual processing, indicating that these regions may be, at least to some extent, predetermined to develop and function. Conversely, the refinement of more complex levels of audiovisual skills appears to require a full multisensory experience throughout lifetime.

The ISC analyses were exploited to evaluate shared representations across visually and acoustically evoked responses in both TD and congenitally SD individuals. Overall, the three experiments revealed a set of regions - STS/STG and neighboring areas - synchronized over time by correspondences between visual and auditory inputs. The congenital absence of visual or auditory experience does not affect the ability of the superior temporal cortex, a brain region devoted to integrated audiovisual processing, to represent unimodal auditory and visual streams pertaining to the same perceptual events. Specifically, high ISC values in temporal and parietal regions were observed for the audiovisual stimulation as a whole. Moreover, when measuring the correlation across the two unimodal conditions, the synchronization was maintained not only in TD participants, but also among the subjects from the two SD groups. However, although synchronized, the ISC magnitude in the congenitally blind and deaf groups was significantly lower than the one measured between conditions within the TD group. In summary, on the one hand, a shared synchronization demonstrates that the ability of these areas to process signals originating from the same natural events in a modality-independent manner is unaffected by the lack of any prior integrated audiovisual experience. On the other hand, the difference between TD and SD indicates that a full functional refinement of audiovisual processing may require the typical interaction between these two senses during development.

In addition to STS/STG, auditory and visual responses also were synchronized within the posteromedial parietal areas (e.g., posterior cingulate and POS) in both TD and SD. On the contrary, synchronization in the left inferior frontal gyrus and in the bilateral medial prefrontal cortex was observed in TD participants only. Moreover, scrambling the chronological order of scenes hindered the synchronization in these parietal regions, providing further evidence for their involvement in discourse and narrative understanding across modalities (see Supplementary Discussion). The same procedure did not impair synchronization in the superior temporal cortex, suggesting that other features are more relevant to this area.

Features modeling was adopted to test the computational properties of synchronized cortical regions. To weigh the relative contribution of distinct stimulus properties, here we developed a novel *model-mediated ISC* approach.

Our low-level perceptual models were inspired by previous studies showing that, during the processing of natural sounds, STS/STG extracts multiple acoustic features, including the amplitude modulation of the temporal envelope^13^, the spectro-temporal modulations^14,15^, the pitch^16^, and the timbre^17^. Moreover, the same regions are also pivotal hubs of the language network^18^. In particular, the middle and posterior portions of STS encode phonetic features^19^, whereas more anterior regions are involved in lexical and semantic processing and contribute to the grammatical or compositional aspects of language^20^.

Regarding the processing of visual properties, there is compelling evidence that STS/STG represents biological motion^21^. Indeed, neurons in middle and posterior STS detect both the snapshots of a biological action and their spatiotemporal integration, as well as complex visual patterns of motion^21^. Taken together, the ability to code static and dynamic properties of visual stimuli, combined with the capacity to process acoustic information and to integrate the two modalities^1^, allows STS/STG to encode multisensory objects^2^, to solve face-voice matching^22^, to represent actions^23,24^ and, more in general, to respond to biologically salient events^1^.

In our study, the ISC of STS/STG was significantly moderated by spectral and amplitude modulations of sounds, and by static Gabor-like filters and motion energy of visual input in both TD and SD individuals. This suggests that low level properties of both auditory and visual inputs exert a crucial role for the computations performed by these regions. This observation is in line with previous evidence showing that the dynamic properties of visual and acoustic signals during speech processing (i.e., lip contour configuration and sound envelope) are correlated and drive stimulus perception as it occurs in the McGurk effect^25^.

Regarding high-order properties, either defined by means of word embeddings or categories, previous evidence suggested the involvement of STS/STG in semantics^26,27^. Other studies proposed that semantic representations may instead retain a large-scale organization distributed throughout the cortex^18,28^. Here, we observed that the high-level movie properties, such as word embeddings and categorical semantic information, mediate the synchronization across auditory and visual stimulations in TD but not in the SD participants. While perceptual features are intrinsic properties of a given stimulus, higher order characteristics rely more on experience and learning, and this may partially account for the observed differences between TD and SD participants. Of note, a significant ability of STS/STG to represent congruent events across the two sensory streams is still present by means of low-level models across samples. However, we cannot determine the extent to which this synchronization is affected by atypical and heterogeneous developmental trajectories that are known to alter language processing distinctively in congenital blindness^29^ and deafness^30^ (see Supplementary Discussion).

Additionally, in order to avoid these coarse correspondences between the two sensory streams, all the models were cleaned from collinearities related to the editing properties of the narrative. Indeed, these features may affect the low-level perceptual (e.g., transitions between scenes would likely correspond to coherent changes in images and sounds) and the high-level semantic descriptors (e.g., spoken, and written dialogues, see Supplementary Results and Discussion).

When measuring the temporal characteristics of the correspondences between visual and acoustic features in the movie, a series of highly coherent events, lasting from tenths of a second to minutes, were found. Such correspondences in the stimulus properties provided the basis for the TRW analysis that estimates the temporal tuning and the duration of receptive windows in the brain. TRW analysis revealed temporal windows ranging from ten seconds (e.g., A4, A5 and STSd) to a couple of minutes (e.g., middle and anterior STSv, PostCing, POS), in line with fMRI temporal resolution. The temporal dynamics of stimulus processing were strikingly consistent across our experimental groups and were arranged to form chronotopic maps, with adjacent patches of cortex showing distinct temporal tunings. These findings are coherent with previous studies that manipulated the temporal structure of visual^10^, auditory^31^, and multisensory^32^ naturalistic stimulations: early primary areas exhibited shortest TRWs, whereas high-order brain areas (e.g., TPOJ, PostCing, POS, precuneus and frontal regions) elaborated and integrated information that accumulated over longer time scales. Equally, when analyzing how events are hierarchically processed at multiple time scales in the brain, high-order areas (including TPOJ and the posterior portions of the medial cortex) represent long-lasting, more abstract and modality-independent events^33^.

As far as the topographical arrangement of temporal features in the STS/STG is concerned, literature mainly focused on the mapping of acoustic (and language-derived acoustic) properties. Specifically, an anterior to posterior gradient was demonstrated when processing phrases, words, and phonemes, respectively^34^. Furthermore, a gradient representing features extracted from the speech envelope, as well as from synthetic stimuli, also was found in the central portion of STG^35^. Specifically, within this central portion of STG, the more anterior part represented slower sounds with relatively high spectral modulations (e.g., syllabic, or prosodic timescales), whereas the more posterior one encoded faster sounds with a relatively low spectral modulation (i.e., phonemic timescale). Our topographical organization supports the observation that encoded information in STS/STG represents acoustic, rather than semantic, properties^35,36^. More importantly, the existence of a coherent topographical organization was demonstrated even for visual-only stimulations, favoring the hypothesis of a modality-independent functional organization in the STS/STG.

Altogether, our TRW analysis in the temporal and parietal areas endorses the possibility of a representation of correspondences across modalities based on a hierarchical temporal tuning, that is maintained despite atypical postnatal trajectories of sensory development.

Individuals born with the congenital lack of a sensory modality have been offering the opportunity to understand the extent to which prior sensory experience is a mandatory prerequisite for the brain organization to develop^7,8,37,38^. Here, two distinct models of sensory deprivation were specifically studied using the same stimulus content, as conveyed through the spared sensory channel. The functional features shared between the two SD models, and then with the TD groups across unimodal and multisensory stimuli, imply that the superior temporal cortex processes congruent signals related to the same event -in a modality-independent manner and autonomously from a prior (multi)sensory experience - and is prevalently predetermined to act as a “correlator” of perceptual information across modalities.

Previous evidence has been showing that the morphological and functional large-scale architecture of the human brain results to be largely preserved in SD models despite the lack of a sensory input since birth and is characterized - to some extent - by modality-invariant computations across several functional domains^7,8,37,38^. The evidence of this supramodal organization has been extended here also to multisensory cortical regions. As a matter of fact, observations in newborns^39^ and sensory-restored individuals^40,41^ already indicated that basic integration of multiple sensory inputs does not require an early multisensory experience to develop^4,42,43^ and that the functional architecture needed for multisensory processing may be already present within the first weeks of life^44^.

Equally, our results also show that audiovisual experience is required for a full refinement of multisensory functions within temporal and parietal regions. Specifically, in the STS/STG both the reduced synchronization and the absent representation of higher-level features in SD individuals suggest that sensory experience during early development is necessary for a complete maturation of functional specializations. In line with this, audiovisual experience appears to be pivotal for the full development of more mature, higher-level computations^5,6^ (e.g., speech perception or semantic congruence).

Altogether, the present findings indicate that human areas subserving audiovisual processing may be innately provided with a functional scaffolding to yield basic multisensory representation and perceive coherence across sensory events. The existence of a ‘proto-organization’ in the multisensory areas is aligned with the morphogenetical and functional evidence that large portions of the human neocortex may possess a predetermined architecture, that is based on a sensory-grounded representation of information and forms the scaffolding for a subsequent, experiencedependent functional specialization^45^. Indeed, the innate presence of a topographic organization of the visual and sensorimotor systems provides the foundations for the progressive development and refinement of vision-^45–48^, somatosensory- and motor-related functions^49^. Equally, previous experimental evidence in SD models nicely matches with this hypothesis of an architecture, characterized by topographical maps already at birth and whose refinement is then favored by the cooperation of distinct sensory inputs^45^.

To conclude, here we studied three distinct samples of TD and two models of congenital SD individuals presented with the same narrative via unimodal or multisensory streams to understand the role of innate presence vs. experience-dependent development of the processing of multisensory (i.e., audiovisual) coherence. The demonstration of a preserved functional topography favors the hypothesis of an innate, modality-independent functional scaffolding to yield basic multisensory processing. Within the old ‘nature versus nurture’ debate on the development of multisensory processing, our study sheds new light on the extent to which audiovisual experience is a mandatory prerequisite for multisensory cortex to develop and become able to detect coherent features across modalities.

## METHODS

### Participants

Fifty subjects took part in the study. We enrolled both typically developed (TD) individuals and sensory deprived (SD) subjects, who lack visual or auditory experience since birth.

Three samples of TD individuals underwent a different experimental condition consisting in the presentation of one version of the same movie: either i) the full multimodal audiovisual (AV) (n=10, 35±13 years, 8 females), ii) the auditory (A) (n=10, 39±17 years, 7 females) or iii) the visual (V) (n=10, 37±15 years, 5 females) one. SD individuals comprising blind (n=11, mean age 46±14 years, 3 females) and deaf (n=9, mean age 24±4, 5 females) participants were presented with the A and V movie conditions respectively. Two blind subjects were removed from the fMRI analysis for excessive head movement (final sample, n=9, mean age 44±14 years, 3 females). All participants were right-handed, as resulted from the scores of the Edinburgh Handedness Inventory. Blind and deaf participants were congenitally deprived with the exception of one deaf subject that reported sensorineural hearing loss before the first year of age and have no history of any psychiatric or neurological disorders. All deaf individuals were proficient in Italian Sign Language (LIS) and did not use hearing aids at the moment of the study. TD subjects reported no hearing impairment, normal or corrected-to-normal vision and no knowledge of the LIS. Only native Italian speakers were selected to be enrolled in the study. Additional information about the deaf and blind samples is provided in the Supplementary Table 2.

Each volunteer was instructed about the nature of the research and gave written informed consent for the participation, in accordance with the guidelines of the institutional board of Turin University Imaging Centre for brain research. The study was approved by the Ethical Committee of the University of Turin (protocol n. 195874, 05/29/19) and conforms to the Declaration of Helsinki.

### Stimulus

Naturalistic stimulation was provided through the presentation of the V, A, and AV versions of the live-action movie *101 Dalmatians* (S. Herek, Great Oaks Entertainment & Walt Disney, 1996). To facilitate subjects’ compliance, a story with a linear plot was selected to make the narrative easy to follow in the context of unimodal presentation. The movie was shortened to make it suitable for a single scanning session. For this purpose, we discarded the scenes whose exclusion does not alter the main narrative thread and we merged the remaining parts together to ensure smooth transitions among cuts while preserving the continuity of narration. The movie was edited to a final version of about 54 minutes that was then split into six runs (~ eight minutes). A fade-in and fade-out period (~ six seconds) was inserted at the beginning and the end of each run. In addition, a scrambled run (eight minutes length) was built by randomly sampling and concatenating the discarded sequences according to the distribution of the duration of movie cuts. This procedure preserved the same low-level features of the original movie while purposely disrupting the narration.

As concerns the auditory version of the stimulus, a voice-over was superimposed over the original movie soundtrack to convey the information carried by the missing visual counterpart. The Italian audio-description of the movie was adapted to our shortened version of the film. Therefore, several parts of the original script were re-written not only to better bridge the gaps we introduced via editing, but also to ensure a satisfactory verbal depiction of those aspects of the visual scenery that are caught by neither characters’ dialogues nor music valence but, still, are essential for the story understanding. The voice-over was performed by professional Italian actor and recorded in a studio insulated from environmental noise and provided with professional hardware (Neumann U87 ai microphone, Universal Audio LA 610 mk2 preamplifier, Apogee Rosetta converter, Apple MacOS) and software (Logic Pro) equipment comprising a set of microphones and filters to manipulate sounds rendering. Then, the voice track was adequately combined with the movie original soundtracks and dialogues. Fade-in and fade-out effects were introduced to smooth the auditory content at the beginning and end of each run to better manage the transitions among the subsequent segments of the film. Music and voice were finally remixed.

We faithfully transcribed the soundscape (e.g., human voices, narrator voice-over, environmental and natural sounds) of the movie into subtitles. Subtitles were written in different styles and colors according to the speaking voice to facilitate speech segmentation and aid understanding. Since line segmentation does not interfere with either reading and story comprehension or image processing^50^, the subtitle pattern was modified in subsequent visual displays upon necessity, appearing in both two-lines and one-line format. Video editing was carried out using iMovie software (10.1.10) on an Apple MacBook Air, whereas for the creation of subtitles, we rely on the open-source crossplatform Aegisub 3.2.2 (http://www.aegisub.org/). In the visual and audiovisual conditions, a small red fixation cross was superimposed at the center of the display, whereas subtitles were shown in the lower section of the screen.

### fMRI experimental design

Before starting the scanning session, participants were asked to rate their general knowledge of the movie plot on a Likert scale ranging from 1 (not at all) to 5 (very well). Participants of each experimental sample were presented with one of the edited versions of the movie (visual, auditory or audiovisual) while undergoing fMRI recordings. Participants were instructed to simply “enjoy the movie”. Structural and functional data acquisition were performed on a single scanning day. After the scanning session, an *ad hoc* two-alternative forced choice questionnaire about the content of the story was administered to assess subject engagement and compliance. Aside, other psychometric scales were administered to participants (see Supplementary materials).

### Stimulation Setup

Audio and visual stimulation were delivered through MR-compatible LCD goggles and headphones (VisualStim Resonance Technology, video resolution 800×600 at 60 Hz, visual field 30°× 22°, 5”, audio 30 dB noise-attenuation, 40 Hz to 40 kHz frequency response). Both goggles and headphones were prescribed irrespectively of the experimental condition and group membership, meaning that each subject wore both devices. The video and audio clips were administered through software package Presentation^®^ 16.5 (Neurobehavioral System, Berkeley, CA, USA -http://www.neurobs.com).

### fMRI data acquisition and preprocessing

Brain activity was recorded with Philips 3T Ingenia scanner equipped with a 32-channel head coil. Functional images were acquired using gradient recall echo planar imaging (GRE-EPI; TR = 2000 ms; TE = 30 ms; FA = 75°; FOV = 240 mm; acquisition matrix (in plane resolution) = 80 × 80; acquisition slice thickness = 3 mm; acquisition voxel size =3×3×3 mm; reconstruction voxel size =3×3×3 mm; 38 sequential axial ascending slices; total volumes 1,614 for the six runs of the movie, plus 256 for the control run). In the same session, three-dimensional high-resolution anatomical image of the brain was also acquired using a magnetization-prepared rapid gradient echo (MPRAGE) sequence (TR =7 ms; TE = 3.2 ms; FA = 9°; FOV= 224, acquisition matrix = 224 × 224; slice thickness = 1mm; voxel size = 1×1×1 mm; 156 sagittal slices).

fMRI data preprocessing was performed following the standard steps with AFNI_17.1.12 software package^51^. First, we removed scanner-related noise correcting the data by spike removal (*3dDespike*). Then, all volumes comprising a run were temporally aligned (*3dTshift*) and successively corrected for head motion using as base the first run (*3dvolreg*). A spatial smoothing with a Gaussian kernel (*3dBlurToFWHM*, 6mm, Full Width at Half Maximum) was applied and then data of each run underwent percentage normalization. Aside, detrending applying Savitzky-Golay filtering (*sgolayfilt*, polynomial order: 3, frame length: 200 timepoints) in MATLAB R2019b (MathWorks Inc., Natick, MA, USA) was performed onto the normalized runs to smooth the corresponding time series and clean them from unwanted trends and outliers. Runs were then concatenated, and multiple regression analysis was performed (*3dDeconvolve*) to remove signals related to head motion parameters and movement spike regressors (frame wise displacement above 0.3). Afterwards, single subject fMRI volumes were nonlinearly (*3dQWarp*) registered to the MNI-192 standard space^52^.

### Computational modeling

We took advantage of computational modeling to extract a collection of movie-related features. Specifically, two sets of low-level features were defined, one extracted from the auditory stream (spectral^26^ and sound envelope^53^ properties to account for frequency- and amplitude-based modulations) and one from the visual stream (set of static Gabor-like filters -GIST^54,55^ and motion energy information based on their spatiotemporal integration^56^). Moreover, a second set of high-level features was modeled based on a manual tagging of natural and artificial categories occurring both in the auditory and visual streams, as well as word embeddings from subtitles using the word2vec algorithm^57^. Finally, a set of features related to the movie editing process (e.g., scene transitions, cuts, dialogues, music and audio descriptions) were manually annotated. A detailed description of these computational models, as well as the parameters used to extract the features for this specific movie were reported in Supplementary Methods.

First, the relationships in the time frequency space between the visual and auditory streams of the stimulus were examined at the maximum available temporal resolution (i.e., frame duration, which was 0.04 s at 25 frames per second). Here, to measure finer co-occurrences in time between the two streams, we relied on fast static visual features extracted from each frame using the GIST model^54^, and the soundscape acoustic spectral properties^26^. Subsequently, since both models retained high dimensionality which prevented the calculation of an overall measure of coherence between them, two representational similarity matrices (RDM) were built by measuring the Euclidean distances of each timepoint in the acoustic and visual feature spaces. As results, these two RDM described the relationship between each movie frame in the visual and auditory domains. To further reduce the dimensionality of the RDMs, the weighted contribution of each frame was evaluated by summing the Euclidean distances with all the other frames. This procedure, which is analogous to the weighted degree of a node in a fully connected graph, generated a time series for each model which resembled the similarity of movie frames across time. Finally, the time series generated from the visual features were compared with the one obtained from the acoustic ones by means of a Continuous Wavelet Transform^58^ (CWT, default parameters with the Morlet wavelet). Results were depicted in Figure 1C and indicated thousands of timepoints showing high coherence, suggesting the visual and acoustic features were continuously entangled representing common events lasting from tenths of a second to several minutes.

Second, in order to clean all stimulus models from a substantial portion of common variance, each model was orthogonalized from the movie-editing descriptors. Indeed, the movie editing features (e.g., film cuts, scene transitions, presence of dialogues, and music) have an impact both on the low-level (e.g., transitions between scenes are often marked by a switch in the movie visual and auditory properties) and the high-level semantic descriptors (e.g., spoken and written dialogues) and could, in principle, mask the impact of the fine-grained computational features, inflating the explained variance of each model. Thus, for every column of each model, a multiple regression was performed to predict computational features by using the movie-editing descriptors as predictors. This procedure generated residuals from the features which became our computational models used in the encoding procedure and for the model-mediated ISC (both described below). Moreover, this procedure computed the portion of unique variance explained by each model discarding a large percentage of shared information (see Supplementary Figure S2, panel B).

Third, since model dimensionality was largely different across computational models (from a few columns of the acoustic model to hundreds in the visual one), an encoding procedure was performed in the multisensory AV condition only, to both verify the goodness of our descriptors and to prune irrelevant features.

Specifically, we first defined an outern cross-validation leave-one-run-out loop in which a run was used as test set, whereas the other five as training set. In the training set, we defined another inner cross-validation leave-one-run-out loop in which a run was used as a validation set to tune the model parameters (i.e., the smallest set of features which provided the best prediction). During the training phase, for each step of the inner loop, a voxel-wise multiple regression was performed to obtain t-stats of the beta coefficients of model descriptors. The t-stats were calculated by averaging the beta coefficients across subjects and by dividing them for their standard errors. The inner cross-validation procedure generated an averaged across folds t-stat for each feature. Within the inner loop, feature selection was performed by considering the prediction performance (R^2^) of reduced models, obtained by increasingly adding the best features (i.e., the ones with the highest t-stats) from one to six. We decided to limit the maximum number of features which could predict a BOLD signal in a voxel to six, as the encoding properties of small patches of cortex (or voxels) seems to rely on a relatively low dimensionality^59,60^. Then, the selected features were used to predict the test set of the outern cross-validation loop, thus, to provide an unbiased estimation of the goodness of fit. Unthresholded R^2^ maps were reported in Supplementary Figure S1 and demonstrated the overall quality of our encoding procedure across all the models.

Finally, for each model, we retained only the best and most frequently used thirteen features across brain voxel to match the dimensionality of our smallest set of descriptors (the acoustic model). The selected acoustic, visual and semantic features, with equal dimensionality and high predictive power, were used on independent data (i.e., model-mediated ISC in the *A vs V* condition, both in TD and SD participants).

In addition, an overall measure of collinearity (R^2^) was estimated between our final set of computational features, before and after the removal of the movie editing feature, thus accounting for the residual collinearities between the models. In detail, each pair of models were compared so that each model acted as a predictor of the other and vice versa. To do so, the multiple regression was combined with a bootstrapping procedure (10,000 iterations) to randomly sample four columns from the predictor model (i.e., four is the size of the movie editing model which retained the lowest dimensionality) and one column from the predicted model. Ultimately, this procedure generated a predicted model which was compared to the original one by means of R^2^ to obtain a final unbiased estimation of collinearity between models of different dimensionality. Note, that after having cleaned the movie-editing features from all the other models, as expected, they still retained some degree of collinearity (up to 5% of total variance) one with the others (Supplementary Figure S2).

### Across-modality Intersubject Correlation (ISC) analysis

The ISC analysis was first performed in the group of subjects exposed to the multimodal (i.e., audiovisual) stimulation. Therefore, for any given voxel included in our grey matter mask, the preprocessed time series of brain activity was extracted, and the average Pearson correlation coefficient (r) was calculated over every possible pair of subjects^9^. Moreover, an across-modality (*A vs V*) ISC analysis was computed through the synchronization of brain activity across individuals presented with a unimodal version of the movie. Notably, this procedure translates in evaluating, voxel by voxel, the correlation in the BOLD activity evoked in the subjects listening to the movie with that elicited in those watching the movie. Therefore, subject pairings were made *across-modalities* matching individuals exposed to different sensory experiences.

To test the statistical significance of the ISC values, a non-parametric permutation test was run by generating surrogate voxel signals splicing the original data in eighteen chunks (three for each run) which were randomly rearranged and eventually time-reversed (1,000 permutations). This procedure allowed to generate a null distribution which shared the same parameters (e.g., mean, standard deviation) of the original data, as well as similar (but not identical) temporal dynamics. In detail, for each voxel, the correlation coefficient was evaluated over every possible pair of subjects, and from this set of coefficients a t-stat was estimated (i.e., dividing the mean r coefficient by its standard error). The same procedure was also carried out with 1,000 surrogate voxel time series, thus obtaining a null distribution of t-stats which provided the one-tail p-value. P-values were estimated using a generalized Pareto distribution to the tail of the permutation distribution^61^. Correction for multiple comparisons was provided by thresholding statistical maps at the 95^th^ percentile (p<0.05, Family-Wise Error -FWE-corrected) of the maximum t distribution from the permutation^62^. Finally, a conjunction analysis in TD participants between the AV and *A vs V* conditions was performed to evaluate the brain regions commonly synchronized across non-deprived groups. This conjunction analysis aimed at identifying and characterizing a core set of brain areas synchronized during the full multisensory experience and that equally shared a common representation across the unimodal processing of auditory and visual information in typically developed participants. The resulting map was subsequently used for the evaluation of the ISC in the *A vs V* condition in the deprived groups, as well as the Temporal Receptive Windows (TRW) and the model-mediated ISC across all experimental conditions.

### Model-mediated ISC

To assess the role of each model, an algorithm based on mediation analysis^63^ was developed. The idea behind mediation analysis relies on the fact that a mediating factor intervenes in the relationship between the independent and the dependent variables. Here, each model was used as a mediating factor during ISC. Specifically, before computing the ISC, the model contribution in the prediction of the BOLD signal was first removed in each subject separately through multiple regression. This procedure generated an ISC value for each model which represented the residual synchronization among subjects independent from our stimulus descriptors. For example, in a voxel that showed high ISC, a model able to predict most of its neural activity would generate a model-mediated ISC close to zero. Thus, the synchronization across subjects would critically depend on the features represented in that model. Conversely, a voxel showing high ISC before and after the mediation analysis would be interpretable as a voxel with an elevated synchronization across subjects, driven by unspecified (at least according to the considered models) activity. As concerns the low-level models, auditory and visual information were removed from the brain activity of those samples presented with the unimodal *V* and *A* condition, respectively, whereas the high-level semantic model, which was inherently multimodal, was removed from both conditions. To obtain a statistical measure on the mediation effects of our computational models, a permutation test was performed generating surrogate voxel signals splicing the original data similarly to the ISC described above. This allowed us to have a null distribution of model-mediated ISC values for each voxel and model. The statistical analysis was performed by evaluating the ‘drop’ (i.e., ISC *minus* model-mediated ISC) and by comparing its intensity with the ones obtained from the null distribution. P-values were estimated using a generalized Pareto distribution to the tail of the permutation distribution^61^ and corrected for multiple comparisons as above (p<0.05, FWE corrected^62^).

### Temporal Receptive Windows (TRW) Analysis

Sensory, perceptual and cognitive processes in the brain rely on the accumulation of information over different time spans. To measure the hierarchical organization of information processing, we computed the ISC over overlapping, binned segments of data. Specifically, during ISC, the Pearson correlation coefficient was calculated after averaging consecutive time points over overlapping rectangular sliding windows ranging from 2 seconds to 240 seconds and moving in steps of 2 seconds. Subsequently, for each voxel, we extracted the width of the window showing the highest synchronization (i.e., highest correlation coefficient) across subjects, named as Temporal Receptive Window (TRW), which is conceptually similar to the approach of Hasson and colleagues^10,33^. To test the statistical significance of the TRW, similarly to ISC, a non-parametric permutation test was performed by evaluating correlation coefficients using surrogate voxel time series (1,000 permutations) using the specific temporal tuning of each voxel. P-values were estimated using a generalized Pareto distribution to the tail of the permutation distribution^61^ and corrected for multiple comparisons as above (p<0.05, FWE corrected^62^).

### Data and code availability

Data and code are available on the osf.io website. Only preprocessed functional data was shared to comply with the European General Data Protection Regulation (GDPR). Raw structural and functional MRI data are available from the corresponding author upon reasonable request.

## Acknowledgments

This work has been supported by a PRIN grant (2017_55TKFE) by the Italian Ministry of University and Research to P.P. Additionally, F.S. was supported by the Frontier Proposal Fellowship (FPF program, 2019) granted by IMT School for Advanced Studies Lucca. A preliminary version of this manuscript has undergone an internal review by Dr. Lotfi Merabet. We acknowledge Dr. Velia Cardin and Dr. Olivier Collignon for insightful discussions on the project.

## Author information

**Affiliations**

**MoMiLab, IMT School for Advanced Studies Lucca, Lucca, Italy**

Francesca Setti, Giacomo Handjaras, Davide Bottari, Luca Cecchetti, Pietro Pietrini & Emiliano Ricciardi

**Department of Translational Research and Advanced Technologies in Medicine and Surgery, University of Pisa, Pisa, Italy**

Andrea Leo

**Department of Psychology, University of Turin, Turin, Italy**

Matteo Diano, Valentina Bruno, Carla Tinti & Francesca Garbarini

## Contributions

F.S., G.H., D.B., A.L., P.P., and E.R. Conceptualization;

F.S., A.L., and G.H. Methodology, Software & Formal Analysis;

F.S., A.L., and M.D. Investigation;

F.S., G.H., F.G., V.B., and C.T. Resources;

F.S., and G.H. Data Curation;

F.S., G.H., D.B, L.C., F.G., P.P., and E.R. interpreted results of experiments;

F.S., G.H., D.B., and E.R. Writing - Original Draft;

All the authors Writing -Review & Editing;

F.S., and G.H. Visualization;

F.G., D.B., L.C., P.P., and E.R. Supervision;

F.G., P.P., and E.R., Project Administration;

F.G., P.P., and E.R., Funding Acquisition.

