## Supplemental Materials for "A modality independent proto-organization of human multisensory areas"

### Table of Contents

Supplementary Results…………..………………...page 2
Supplementary Discussion………….…………………...page 3
Supplementary Methods…………….………………...page 8
Figure S1…………...……………………………...page 16
Figure S2…………….…………..………………..page 17
Figure S3…………….…………..………………..page 18
Supplementary Table1…………..…………………page 20
Supplementary Table2…………..…………………page 22
Supplementary Table3…………..…………………page 23
References……………………………………………page 28

### Supplementary Results

#### Behavioral assessment

Familiarity with the movie plot was assessed through a Likert scale ranging from 1 to 5. The majority of the participants reported to have a general knowledge of the main facts of the narrative (TD median: 3; SD median 2,5; overall four subjects declared no familiarity at all with the movie, whereas none of them stated to know the story very well).
After the scanning session, an *ad hoc* two-alternative forced choice questionnaire about the content of the story was administered to assess subjects' engagement and compliance. In the first two experimental sessions, all TD participants attended the movie as resulted from our assessment questionnaire (n=30, mean accuracy ± standard deviation: 87% ± 7%; range: min-max: 72-100%). Similarly, in the third experiment, all SD participants attended the movie as resulted from the final questionnaire (n=18, mean accuracy ± standard deviation: 82% ± 13%; range: min - max: 56% - 100%).

### Supplementary Discussion

#### The posterior cingulate cortex is synchronized across meaningful auditory and visual movie

Results of the ISC analysis showed a shared recruitment of the posterior cingulate cortex (a hub of the Default Mode Network -DMN-) across the two unimodal conditions of stimulus presentation (i.e., *A vs V*) that may be associated with high-level cognitive processes related to narrative understanding.

Such finding is compatible with previous evidence reporting a common engagement of the DMN and the STS/STG, during the encoding of naturalistic information (Chen et al., 2017; Gould Van Praag et al., 2017), processing of spoken and written natural language (Regev et al., 2018) and comprehension of naturalistic narrative speech (Dikker et al., 2014; Ferstl et al., 2008; Honey et al., 2012). This hypothesis is further supported by the findings of the current study. First, the results of the model-mediated ISC revealed that the shared processing of visual and auditory streams in the posterior cingulate cortex is mainly driven by high-level features associated to the movie semantics (i.e., linguistic and category-related information). Second, the synchronization across auditory and visual streams in this region is disrupted by the presentation of a scrambled movie condition, in which the chronological order of cuts is purposely altered to make the storyline nonsensical. To this regard, while synchronization of brain activity among viewers does occur in a network of brain areas comprising the posterior cingulate cortex for the processing of both coherent silent videos and meaningful auditory movies, this same synchronization is instead dampened by the presentation of random/scrambled visual sequences or meaningless audio clips (Loiotile et al., 2019). Finally, TRW analysis demonstrated that the temporal dynamics of input processing in this region are relatively slow (of the order of minutes), thus reflecting a high-level stage of information processing, compatible with a role of the posterior cingulate cortex in story understanding. Taken together, these observations suggest that the posterior cingulate cortex may represent the stimulus narrative (Regev et al., 2013) in which meaningful events are accumulated and integrated over relatively long timescales (Ames et al., 2015; Simony et al., 2016).

#### The inferior frontal cortex is not synchronized across SD samples

Integration of signals across modalities relies both on object knowledge and previous experience in relating sensory properties belonging to the same stimulus (Noppeney et al., 2003). Such process counts on the recruitment of a fronto-temporal network comprising perisylvian temporal and inferior frontal regions, prefrontal cortex and the precuneus (Ferstl et al., 2008) which are involved in object recognition (Amedi et al., 2005), comprehension of non-linguistic conceptual information (Koelsch et al., 2004), discourse understanding and working-memory (Baldassano et al., 2017; Santi & Grodzinsky, 2007). Specifically, the left inferior frontal gyrus is engaged during the processing of everyday, bimodal audiovisual stimulation, thus providing a modality-independent neural substrate crucial for relating information that is shared across the two senses (Porada et al., 2021).

For what concerns congenital blindness and deafness, semantic knowledge is acquired both via a sensory-grounded event representation dependent on the spared senses (Amedi et al., 2007; Ricciardi et al., 2014), and through high-level, sensory independent mechanisms based on linguistic description and cognitive inference (Handjaras et al., 2016; Wang et al., 2020) and relying on a distributed network of regions comprising the frontal cortex. Additionally, large-scale modifications in the functional coupling among brain areas has been demonstrated in congenitally blind individuals which present an enhanced functional connectivity between the primary visual cortex and the inferior frontal gyrus/sulcus (Bedny, 2017; Deen et al., 2015; Noppeney et al., 2003).

Moreover, ISC analysis revealed congenitally blind individuals commonly engage the inferior frontal gyrus/sulcus and middle frontal junction while listening to auditory movies (Loiotile et al., 2019). Similarly, a recruitment of the left inferior/middle frontal gyri has been found in congenitally deaf signers during the presentation of isolated written English words and pictorial stimuli (Waters et al., 2007). Therefore, we expected to find a significant ISC in the inferior frontal gyrus across visual and auditory movie processing in both TD and SD samples. However, we found that only in TD participants the left inferior frontal gyrus and the bilateral medial prefrontal cortex were synchronized across auditory and visual conditions, while synchronization across deaf and blind participants did not occur. Several explanations can account for this finding. To begin with, this lack of synchronization may depend on the differences in language processing between the two groups of congenitally deprived participants, whose post-natal sensory experiences inevitably differ. Indeed, the absence of specific sensory experiences early in life affects the development and the (left) lateralization of the neural systems devoted to language processing: while an overlapping left-lateralized, fronto-temporal circuitry is engaged during the processing of information in the native language (for a meta-analytic review see Trettenbrein et al., 2021) both for native signers and hearing individuals during speech, when asked to read written sentences, native signers do not show activation in these areas (Neville et al., 1998). Therefore, given this evidence and considering the properties of our stimulus that requires subtitles reading, we can speculate that our native signers minimally recruited the left frontal regions while watching the movie. Analogously, congenital loss of visual input in blindness affects language processing and determines a reduced left-lateralization of language functions (Lane et al., 2017; Pant et al., 2020). Consistently, we previously showed that the left

IFG was not included in cortical regions involved in semantic processing across sighted and congenitally blind individuals while performing a property-generation task with concrete nouns, presented through visual and/or auditory modalities (Handjaras et al., 2016). Differences in the neural correlates of naturalistic information processing across sensory deprived groups could indeed emerge as a result of differences in visually- and acoustically- based semantic representation of events. In fact, evidence demonstrated that the left inferior frontal gyrus is involved in semantic processing of sentences (Friederici et al., 2003; Hagoort et al., 2004) and is sensitive to the semantic congruence across stimuli, as conveyed by different sensory modalities such as pictures-sounds (Hein et al., 2007) and actions-language (gestures and pantomimes) combinations (Willems et al., 2009). Moreover, evidence exists of a disconnection between sensory regions and higher order areas in sensory deprivation (Kral et al., 2017) and this may possibly explain the lack of synchronization of IFG cortex across blind and deaf individuals. Thus, left IFG participates in the online matching of information across two sensory streams (e.g., gestures with the accompanying speech) to build a novel, context-dependent semantic representation of the external events (Hagoort, 2005). Given these considerations, we can assume that the lack of multisensory experience in coupling speech sounds with the related visual gestures in both deaf and blind participants may prevent the creation of common, shared representations of the same sensory events.

**Movie editing drives ISC across conditions**

A movie results from the editing of quick shots into scenes: artificial sequences that retain the unity of time and location through an accurate mapping of the auditory soundscape with the visual stream. Such a technical process determines the pace of the audiovisual stimulation and the final rhythm of the narrative, ultimately creating a definite stream of meaningful sequences (i.e., scene transitions and camera cuts) that convey the story. Since synchronization of brain activity in fMRI mainly arose from slow-frequency fluctuations (Hasson et al., 2004), we explored the structure of the movie in order to model what we called the “movie editing” features. Therefore, with this term, we refer both to the editor choices (e.g., scenes, camera cuts, dialogues, soundtracks) and to the modifications we researchers made, namely the inclusion of the audiodescriptions and subtitles.

Note that the movie editing model, by the way it is conceptualized, can be conceived as a ‘blanket’ term, encompassing a set of non-specific, visual, auditory and linguistic stimulus properties that cannot be unequivocally assigned to any of the other stimulus models we included in our analysis pipeline. In fact, changes in the movie scenery, for instance, likely result in modifications of the visual properties of the image, often accompanied by variations in the soundscape. Therefore, not surprisingly, it shared a consistent portion of variance in the data with all the other computational models we included in our analysis (Supplementary Figure S2), thus representing an ideal covariate to control for slow temporal changes which might be modulated by other brain processes (e.g., working memory, attentional mechanisms, change in arousal, etc.) apart from sensory processing (Hasson et al., 2015).

Results of the encoding of the editing model (Supplementary Figure S2), revealed that these features significantly impacted brain synchronization across the whole cortex, ranging from the visual and auditory regions, as well as ventral and dorsal attention networks in the frontal and parietal lobes (Andersen et al., 2006). This observation suggests that the movie editing descriptor captures some slow-changing stimulus characteristics, mediated both by bottom-up and top-down (e.g., attentional, arousal and memory) modulations. Such hypothesis follows from the idea that movie understanding requires concurrent deployment of several perceptual and cognitive processes necessary for the creation of a meaningful and coherent representation of the information impinging our senses.

**Limitations**

Our study presents the following limitations.

First, since the aim of the present study was to assess whether the emergence of audiovisual processing in the human brain requires audiovisual experience, the work mainly focused on the study of synchronized brain responses across experimental groups and conditions. Therefore, the investigation of group-specific responses or experience-dependent rearrangements that take place when a sensory modality is absent goes beyond the scope of the current study.

Second, the small sample size (n=18) of sensory deprived groups could result as limited. However, a review of the current literature in the field of sensory deprivation reveals that many studies rely on a comparable number of subjects (e.g., Bedny et al., 2011; Collignon et al., 2011; Watkins et al., 2013) mainly for the difficulties related to the recruitment of these exceptional participants who lack sensory input since birth. For what concerns the field of naturalistic stimulation and ISC analysis, although some studies report bigger sample sizes, they do often rely on the presentation of shorter clips/audio descriptions relative to the long movie (~1h) stimulation adopted here. For this reason, the data of the present work are still aligned with the average number of datapoints reported in the literature (for further details on this please see Supplementary Figure S3 and Supplementary Table 3).

Third, we acknowledge that deaf participants were not matched for age to the other experimental groups, being on average, younger than the other samples (i.e., deaf: n=9, mean age 24±4years; blind: n=9, mean age 44±14 years; TD: AV condition, n=10, 35±13 years; n=10, A condition, 39±17 years; V condition, n=10, 37±15 years).

Even if this discrepancy is mainly related to the exceptionality of congenitally deprived samples and in the difficulty of their recruitment, which does not allow a ‘selection’ based on age, a parsimonious evaluation of this age gaps between samples could not justify the differences that has been reported between groups. In addition, here, deaf individuals have been primarily analyzed for their commonalities with the blind group and the TD samples.

Lastly, we have to mention that subtitles and lip-movement in the visual movie are not fully congruent, since they rely on Italian for subtitles and English for acting, as commonly occurs in Italy for dubbed movies. Nevertheless, even if not specifically determined with an eye-tracking measure, we believe that the presence of subtitles hardly allowed participants to concurrently perform lip-reading. Furthermore, this discrepancy would solely affect the V-only version of the movie for both TD and deaf samples, and -in case- would negatively affect the overlapping responses across samples. Since we are yet demonstrating significant overlapping responses across samples, this point represents an issue of way lesser concern.

### Supplementary Methods

#### Behavioral assessment

The following psychometric scales were administered to participants: one concerning manual dexterity, assessed through the Edinburgh Handedness Inventory (Oldfield, 1971) and a set of questionnaires related to sensory imagery: the shortened version of the Bett's Questionnaire upon Mental Imagery (Sheehan, 1967) the Visual Vividness Imagery Questionnaire (VVIQ) (Marks, 1973) and the Plymouth Sensory Imagery Questionnaire (Andrade et al., 2014).

### Computational models

The following sections will review the models that were adopted in the present work to extract the movie (visual and auditory) low-level and high-level (semantic and categorical) properties. Consistently with the theoretical framework of hierarchical sensory processing (de Heer et al., 2017; DiCarlo et al., 2012; Heeger et al., 1996), we exploited the richness of the naturalistic stimulation to investigate stimulus-driven brain responses to low-level, high-level and categorial movie features. Therefore, a set of computational models of early visual and auditory computations were adopted to extract the frequential signal properties and their modulation in time of both the visual and auditory movie stimuli: features generated from image GIST and motion energy defined the visual model; sound power spectrum and envelope, the acoustic model. Additionally, a semantic high-level model was employed by combining information from the stimulus semantics (e.g., word2vec algorithm - Mikolov et al., 2013) as well as the manual tagging of the categorical content of the visual and auditory movie for event discrimination (i.e., Animals, Houses, Objects, Person, and Vehicles -for the visual stimulus and the auditory track). Finally, a movie-editing model was defined on the stimulus properties (i.e., Cuts, Scenes, Dialogues, Audio Descriptions, Soundtracks), introduced during the editing phase which captured coarse slow-paced collinearities between auditory and visual streams, theoretically having an impact on both low-level and high-level semantic descriptors.

#### Low-level visual model: GIST feature space

A scene GIST model (Oliva & Torralba, 2006) was used to quantify the spatial properties of the movie frames convolving a set of Gabor-like filters with a specific frequency and orientation to the image. Each movie frame was segmented into a 4x4 grid and the responses to Gabor filters having four different sizes and four orientations was sampled, resulting in a model comprising 256 features for each frame (as in Lettieri et al., 2019). Each feature represented the total energy at a particular orientation and spatial frequency, averaged over a position of the visual field. Subsequently, GIST descriptors across 50 frames within 2 seconds were averaged to match the temporal resolution of fMRI. Subsequently, descriptors were normalized, and a Principal Component Analysis (PCA) was applied to retained components able to explain at least 90% of the total variance, thus reducing the model to 22 dimensions. Finally, the remaining columns were convolved with a standard gamma function as the hemodynamic response function.

#### Low-level visual model: Motion energy feature space

The total motion energy was computed for each movie second through a set of 4,715 motion energy descriptors consisting of a quadrature-pair of space-time Gabor filters (e.g., Gabor wavelets with three different temporal frequencies at 0 -static energy-, 2, and 4 Hz as in (Nishimoto et al., 2011).
MATLAB code is available here: https://github.com/gallantlab/motion_energy_matlab.
The model described each movie frame by a set of preferred spatial frequencies, orientations and temporal frequencies that grasp fast-changing visual information. Subsequently, descriptors were normalized and a PCA was applied to retained components able to explain at least 90% of the total variance, thus reducing the model to 398 dimensions. Finally, the remaining columns were convolved with a standard gamma function as the hemodynamic response function.

#### Low-level auditory model: Power Spectrum feature space

Spectral features extraction was carried out following the method described by de Heer and colleagues (2017). We estimated the signal power spectrum for each run through the Welch's power spectral density estimate (Welch, 1967) with a Gaussian window (SD of 5 ms, length 30 ms, 1 ms spacing between window) over portions of the signal of 2 seconds length (in order to match the fMRI temporal resolution). The output is a 449-dimensional vector that summarizes the signal power spectrum (expressed in dB units) in the range of 0 Hz to ~15000 Hz computed in bands of 33.5 Hz. For further details about the parameters used please refer to de Heer et al., 2017; Lettieri et al., 2019. Subsequently, descriptors were normalized and a PCA was applied to retained components able to explain at least 90% of the total variance, thus reducing the model to 5 dimensions. Finally, the remaining columns were convolved with a standard gamma function as the hemodynamic response function.

### Low-level auditory model: Envelope feature space

To model sound amplitude changes over time, the soundtrack envelope power spectral density was extracted. First, we first evaluated the upper and lower root-mean-square envelopes of the raw sound signal averaged over the two channels, through the MATLAB function *envelope* (option '*rms*') with a sliding window of 10 ms length. The signal power spectrum was estimated over 2 seconds signal bins with the Welch's power spectral density estimate (Gaussian window, SD 800 ms, length 1 s, 0.5 s spacing between windows). The output is a 49-dimensional vector that summarizes the envelope power spectrum (expressed in dB units) in the range of 1 Hz to 99 Hz computed in bands of 2 Hz. For further details, please refer to Martinelli et al., 2020. Subsequently, descriptors were normalized and a PCA was applied to retained components able to explain at least 90% of the total variance, thus reducing the model to 8 dimensions. Finally, the remaining columns were convolved with a standard gamma function as the hemodynamic response function.

Therefore, the visual model was represented by a matrix of 1,614 rows (as the number of timepoints of the fMRI) and 420 columns (i.e., features), whereas the acoustic model comprised 1,614 rows and 13 columns.

### High-level model: Computational semantic feature space

Given the transcript of the whole verbal content of the movie, comprising all the spoken parts (dialogues, monologues, narrator voice and even animal sounds), we manually chucked it word by word and got rid of articles, prepositions and pronouns. Note that only nouns, verbs, adverbs and onomatopoeic words were spared and then used to derive the semantic representation of the story. We decided to include also onomatopoeic paralinguistic utterances because they directly mimic specific non-speech and non-musical sounds (produced by nature, animals or human activities), whose source is easily recognizable and inherently *signify* what they refer to. The semantic representation of each term in the transcript was derived through word2vec, a word embedding technique (Mikolov et al., 2013) and based on the distributional properties of words in a large corpus of text. Thus, to identify the movie semantic features, we used the itWaC corpus (Baroni et al., 2009). The itWaC corpus consisted of 2 billion Italian words extracted from the Web (http://wacky.sslmit.unibo.it/doku.php?id=corpora), and it was the largest Italian corpus currently available. Therefore, we used the *word2vec* algorithm to obtain the embedding space (128 dimensions, window 5, cbow architecture, pre-trained in Dell'Orletta &

Nissim, 2018). As a result, we extracted for each 2 seconds interval (fMRI temporal resolution) a 128-sized vector obtained by averaging all the word vectors included in the time frame.  Subsequently, descriptors were normalized and a PCA was applied to retained components able to explain at least 90% of the total variance, thus reducing the model to 72 dimensions. Finally, the remaining columns were convolved with a standard gamma function as the hemodynamic response function.

### High-level model: Categorial features space

Category selective regions in the brain are known to be tuned for processing specific classes of stimuli (Epstein, 2008; Gould Van Praag et al., 2017; Kanwisher & Yovel, 2006; Martin, 2007; McCandliss et al., 2003; Peelen & Downing, 2017). Here, we evaluated the specific contribution of non-linguistic, high-level semantic information in modulating brain activity. We relied on previous literature on the topic (Grill-Spector & Weiner, 2014) for the definition of the visual categories that, after being validated through comparison with the image segmentation output of an automatic algorithm (see below Visual categorial model validation), were used to label the categories of the auditory track as well.

For the visual condition the continuous stream of information was classified in seven categories: *Animals, Body-parts, Faces, Houses, Objects, Persons, and Vehicles*. The very same tagging procedure was applied to the auditory stimulus as well but instead of using all the seven visual categories, we used the notation based on five major classes derived from the validation procedure. Note that, the two stimulation conditions intrinsically vary in the degree of detail conveyed by the diverse sensory modalities: although the presence of the narrator is meant to describe visual information through speech, the acoustic content still communicates information at a broader scale than the visual one. Think about a dialogue between two characters: in the visual condition is possible to appreciate either the *Face* appearance, the *Person* silhouette, posture, clothing or a specific *Body-part* while in the auditory setting we can just say, globally speaking, that a *Person* is present (a general idea/mental representation of a man or a woman which appearance is up to the listener). These modality-dependent aspects and the need for consistency in the methods across stimulus conditions motivated the choice of the following five auditory categories: *Animals, Houses, Objects, Persons, and Vehicles*. Everyday life hearing depends on the selection of informative sounds among less-relevant background noise. Therefore, our classification was mainly focused on non-stationary foreground sounds, namely those sounds whose signal statistics change over time and result to be more informative of the world around us. However, previous work showed that the presence of background environmental “noise” differentially affects primary and non-primary auditory areas responses to concurrent foreground sounds (Kell & McDermott, 2019). Additionally, we can assume that background natural sounds represent a reliable source of information for blind individuals. For these reasons, the classification was also extended to include audio signals from nature, animals, man-made objects and human activities (Mattioni et al., 2020).

**Annotation procedure.** One of the authors of this study carried out the tagging procedure. The stimulus was explored run by run: the relevant features were manually annotated along with their timecode (time resolution of one second).

**Visual categories.** For each time window of 1 second, the annotator wrote down the elements located in foreground and when present, also additional items that for their appearance (color, size, change in position) or story relevance (key information, main characters) capture the viewer's attention. Each entry was then properly classified according to the following seven categories.

*Animals:* all species of animals represented in the movie.
*Body-parts:* it applies only to a given part of the body appearing in isolation and foreground, except from faces. A leg, toe or hand are examples of body-parts.
*Faces*: all the faces represented in foreground regardless of viewpoint and lighting. Faces visible from distance and in presence of the trunk or the entire body are not considered falling in this category (see person).
*Houses*: it refers both to a façade of a building in isolation and to groups of edifices or cityscapes. It also comprises other kinds of structures that are not “houses” in a narrow sense but still, pertains to the more general concept of “buildings” (e.g., a farm, a castle).
*Objects*: it collects man-made objects and tools.
*Persons:* it includes the images in which the complete silhouette of the body is visible, or the head and the upper body. Faces in isolation are not falling into this category.
*Vehicles:* this category is meant to include all “means of transportation” represented in the movie (e.g., a car, a bicycle, a truck) or parts of them sufficiently big and detailed to be recognized as pertaining to a specific kind of vehicle (e.g., a bicycle handlebar or tire; a car hood).
Subsequently, visual categorial descriptors were resampled to match the temporal resolution of fMRI and normalized. Then a PCA was applied to retained components able to explain at least 90% of the total variance, thus reducing the model to 5 dimensions. Finally, the remaining columns were convolved with a standard gamma function as the hemodynamic response function.

**Visual categorial model validation.** To validate the quality of the categorial tagging, we ran an automatic labeling of the content of the visual scenery and tested the degree of classification similarity between the two methods. We relied on a specific kind of a pre-trained convolutional neural network (CRF-RNN) (Zheng et al., 2015) to solve image segmentation and classify the elements of the visual display into five categories (*Animals, Houses, Objects, Person, Vehicles)*. In this approach, pixel-level labels are predicted combining the strengths of Convolutional Neural Networks (CNNs) technique with Conditional Random Fields (CRFs)-based probabilistic graphical modeling. The model comprises two stages: an initial full convolutional deep network followed by a CRF-RNN step, that can be effectively used to accomplish categorial image segmentation tasks. Since the neural net labels slightly differ from the classes that we manually labelled, some categories were grouped together to compare the two approaches. Therefore, aggregated person, faces and body-parts were aggregated in a single descriptor called *whole-person* to adjust for the absence of such finer distinctions in the classes provided within the CRF-RNN (where *person* refers to all the previous). On the other hand, we clustered the net labels referring to the same superordinate category (i.e., airplane, bicycle, boat, bus, car, motorbike, train into *vehicles*; bird, cat, cow, dog, horse, sheep into *animals*; bottle, chair, dining table, potted plant, sofa, tv monitor into *objects*). The rank correlation (Spearman's ρ) was computed to assess reliability across the two classification methods: *animals* (0.518, CI 95: 0.476 - 0.559), *objects* (0.153, CI 95: 0.098 - 0.211), *vehicles* (0.384, CI 95: 0.321 - 0.442); *persons* (0.616, CI 95: 0.580 - 0.650). Discrepancies in the classification among the two procedures were carefully reviewed by two authors of the present work who double-checked and assessed the goodness of the manual tagging.

**Auditory categories.** The categorial content of the auditory movie was described following the same method that we applied for the visual stimuli. Therefore, the auditory track of each run was sampled at the time resolution of one second and the salient sounds were labeled based on the category they pertain to. Aiming for consistency across models, we adopted the same categories that were used for the labeling of the visual movie after the validation step. Therefore, each sound was classified according to one of the following labels.

*Animals:* all kinds of animal sounds audible in foreground or still clearly recognizable from the background audio track.
*Houses*: it refers to the descriptions of houses, buildings or cityscape appearance. Usually made by the narrator's voice, such portrayals make explicit reference to the

presence of a building (not necessarily “houses” strictly speaking). Examples are: “in front of Anita’s house”, “outside, from the castle gate”.
*Objects:* it collects overt reference to objects or refers to sounds that are generated by man-made objects or tools (e.g., the bell ringing, the shower, the teacups chinking).
*Persons:* the presence of a person is mainly denoted by speech or dialogues. Moreover, this category applies to descriptions of a person's appearance and to those sounds that can be unmistakably attributed to a human being (e.g., footsteps, cough, background chattering, screaming).
*Vehicles:* this category includes vehicles' sounds and descriptions of their appearance. All the onomatopoeic sounds recalling vehicles are included in this category (e.g., “wrooooom”, “beep”, “slam”, “screech”).
Subsequently, auditory categorial descriptors were resampled to match the temporal resolution of fMRI and normalized. Then a PCA was applied to retained components able to explain at least 90% of the total variance, thus reducing the model to 4 dimensions. Finally, the remaining columns were convolved with a standard gamma function as the hemodynamic response function.

Therefore, the high-level semantic model was represented by a matrix of 1,614 rows (as the number of timepoints of the fMRI) and 81 columns.
All three models (i.e., auditory, visual, semantic) were also used in an encoding procedure to predict brain activity of fMRI data from the multisensory AV condition. This step tested the overall quality of the models (Supplementary Figure S1), as well as it gave us the opportunity to perform a feature selection procedure, further reducing the dimensionality of each of them to a small set of predictors (i.e., 13; please refer to the *Methods* section, *Computational modeling* paragraph, in the main manuscript).

#### Movie-editing model

Movies are complex stimuli not only for the multifaceted information they convey but also because of their formal architecture, as it results from the work of the film editor. Indeed, the stylistic choices (e.g., camera’s cuts selection, scenes arrangement and duration) build up the peculiar features of the movie framework that, possibly, influence brain activity (e.g., a change in the scene setting may correspond to modifications in the image luminance and be associated with corresponding adjustments in the music). Moreover, we contributed to the process of editing as well, shortening the original duration and modifying both the auditory and the visual streams. To investigate whether these formal aspects influence movie perception, rather than focusing on the content of the stimulus, we explored the structure of the stimulus in order to model what we called the *movie-editing* features. With this term, we thus refer not only to the editor choices already present in the original version, but also to the major modifications that we introduced, namely the inclusion of the audio descriptions and subtitles. We modeled the movie editing features using the same approach devised for the category: four visual (i.e., cuts, scenes, subtitles and text embedded in the visual frames) and three auditory (i.e., audio descriptions, dialogues and soundtracks) movie-editing classes were binary tagged in a window lasting 1 second, as reported below.

*Cuts*: this term is used to define sudden changes in camera angle, location and placement from one shot to the following. These events occur frequently during the narration and can be easily spotted in a glance.
*Scenes*: this label refers to the major changes in the story setting (location, characters, actions and time). Thus, we considered a scene as a story unit that takes place in a specific location and in a defined period of time. Compared with cuts, these events happen on a slower timescale.
*Subtitles:* it reflects the story script including all the spoken parts: voice-over, dialogues and environmental sound (mostly animal sounds).
*Text:* all written text readable on the screen *and* belonging to the original movie (it does not include the subtitles that we added afterwards to the visual version of the movie).
*Audio descriptions:* all the parts of the movie script reported by the narrator voice-over. These descriptions are meant to convey the salient aspects of the tale that cannot be inferred merely by listening to the original movie auditory track because they are generally conveyed by the visual scenery. Indeed, audio descriptions contain mostly scenes portrayals for better contextualization and characters actions/emotional state depiction. Note that this category does not include dialogues, environmental sounds and music.
*Dialogues:* it refers only to the part of the discourse pronounced by a person. This includes both conversations, and “monologues” (e.g., the reporter announcing the news to the spectators, the priest speaking to the audience in the church).
*Soundtracks:* background music tracks.

Subsequently, movie-editing descriptors were resampled to match the temporal resolution of fMRI and normalized. Then a PCA was applied to the retained components able to explain at least 90% of the total variance, thus reducing the model to four dimensions. Finally, the remaining columns were convolved with a standard gamma function as the hemodynamic response function.
The goodness of this model was tested in an encoding procedure to predict brain activity of fMRI data from the multisensory AV condition (Supplementary Figure S2, panel A). Moreover, we measured the collinearity between this model and all the others Supplementary Figure S2, panel B). Since movie-editing features share a large portion

of variance across all the other models, we decided to orthogonalize all the features for the stimulus properties related to the film editing process.

### Supplementary Figures

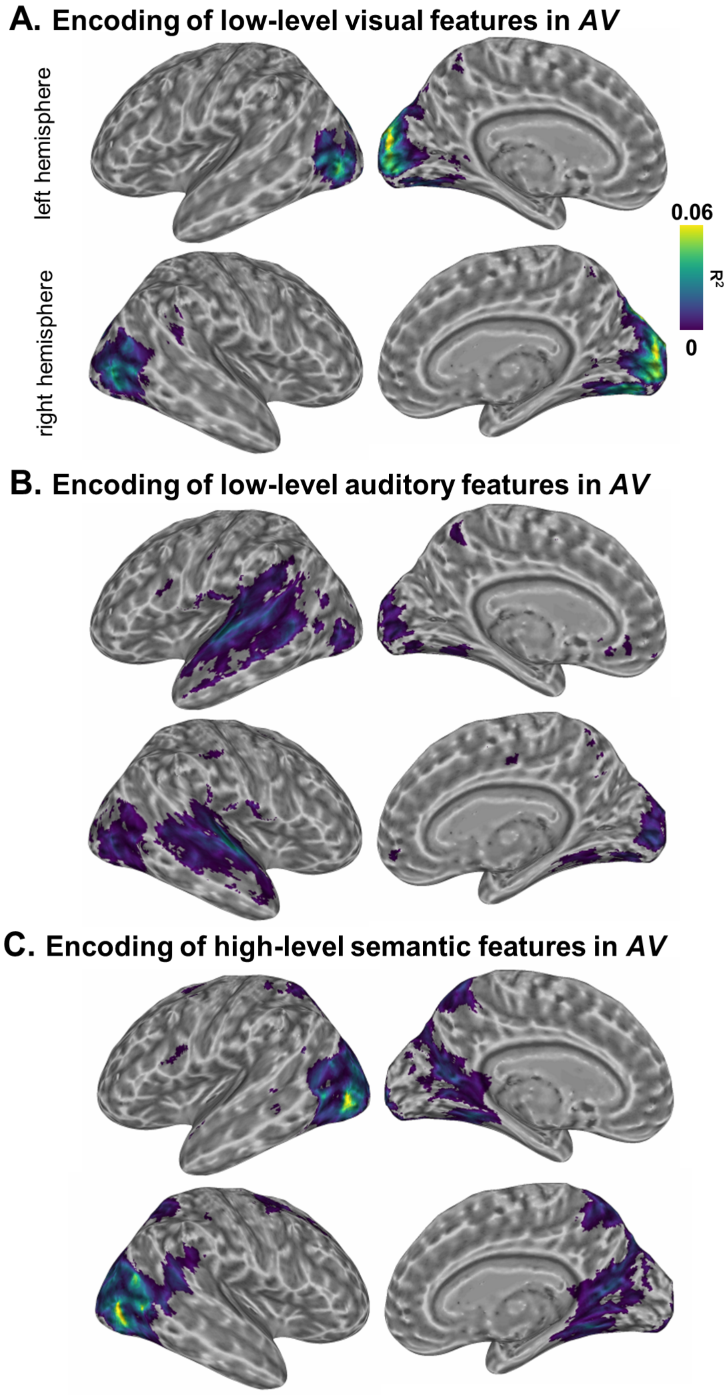


**Figure S1.** Panels A, B, C depict the encoding we performed on the AV condition, for the low-level visual, the low-level acoustic and the high-level semantic features respectively. Unthresholded cross-validated $R^2$ maps, averaged across subjects, show the quality of the encoding procedure for different computational models.

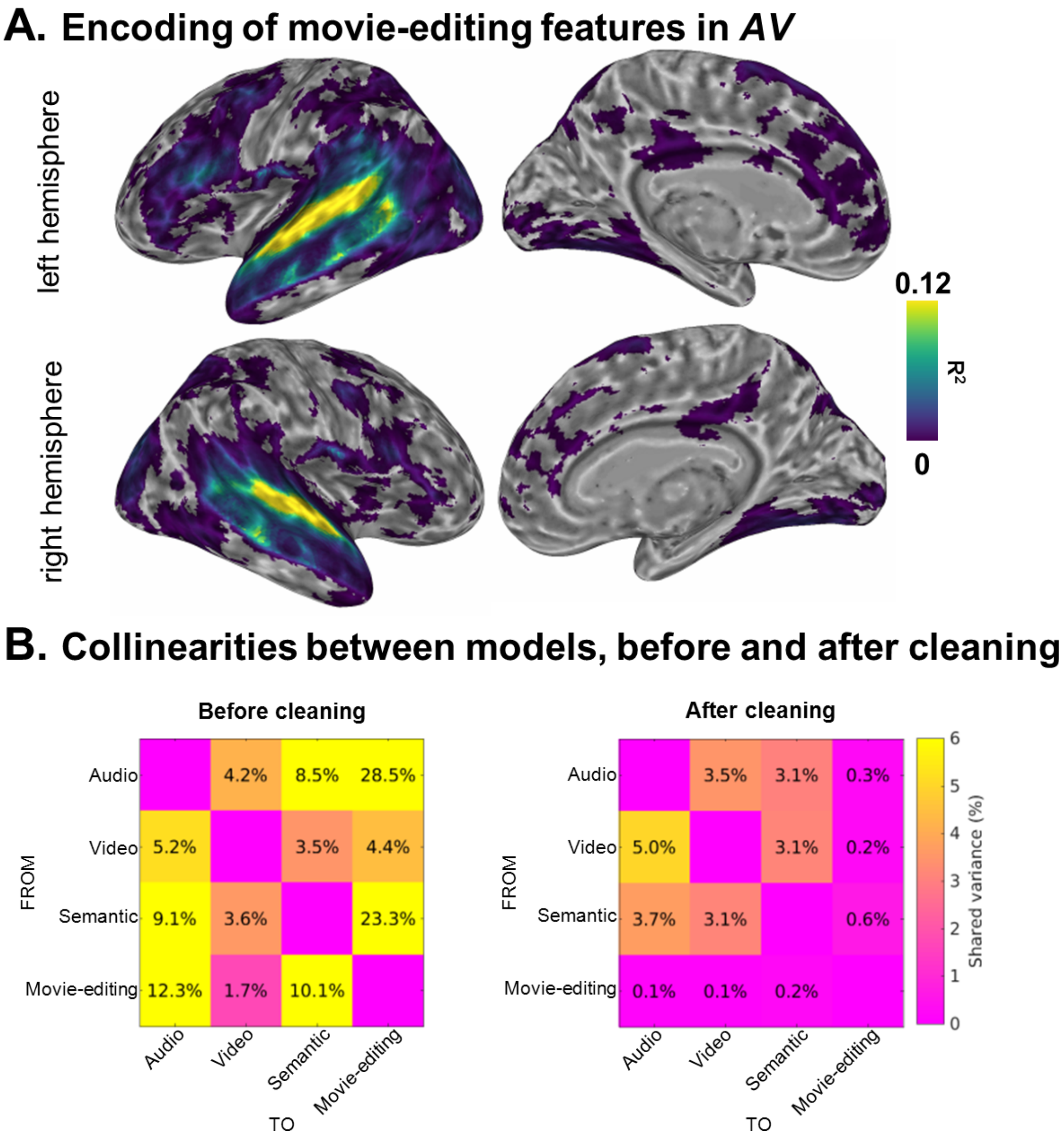


**Figure S2.** Panel A shows the unthresholded cross-validated $R^2$ map, averaged across-subjects, for the encoding of the movie-editing features during multisensory AV stimulation. The matrix on the left in Panel B demonstrates that our set of stimulus features are collinear and therefore share a large percentage of variance. Since both low-level and high-level models are affected by the coarse properties related to the movie-editing descriptor, each of the above was orthogonalized by the latter with the aim to clean them from the portion of common variance. Residual collinearities across models after cleaning are shown in the matrix on the right side of Panel B.

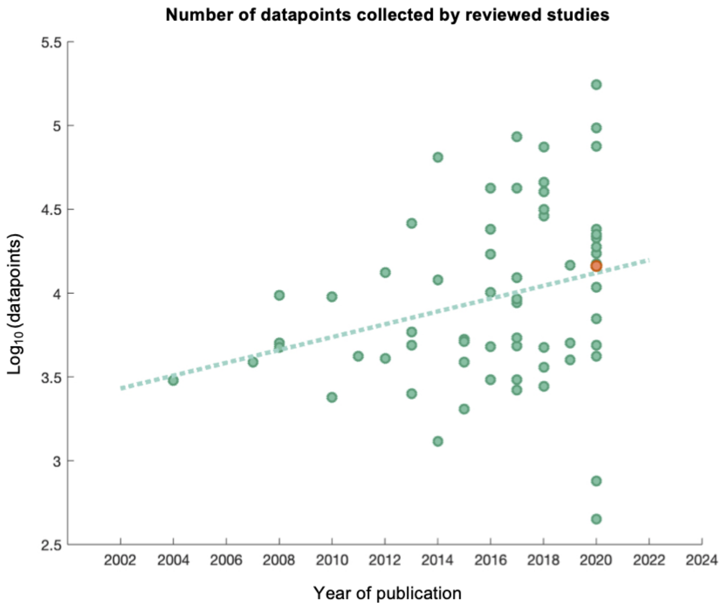


**Figure S3.** Number of acquired data points as a function of the year of publication across fMRI studies using ISC. In the scatter plot, each green point refers to a reviewed paper while the orange dot identifies the study presented here. Data are plotted as the number of functional scans multiplied by the number of subjects on a logarithmic (base 10) scale. The number of data points collected in our study align with that of other works relative to year of publication.

Studies were retrieved after querying on PubMed and Google Scholar in September 2020 the keywords: “fMRI”, “ISC”, “naturalistic stimulation”, “movie”, “narratives” and “stories”. Each time a paper matched our search criteria we processed it and we collected the following data (see Table 3): i) authors and year of publication; ii) name of the movie/track (when available); iii) experimental conditions (i.e., auditory, visual, audiovisual); iv) stimulus length (in seconds); v) sample size; vi) fMRI parameters: TR and number of volumes acquired (when specified). This information allowed us to calculate the amount of data points collected in each study, referring only to the sample of interest. When the same dataset was adopted in multiple publications, we decided to report the year of the work published first. In the case of studies performing ISC analysis on previously acquired datasets, we scrutinized the original study and considered it only in the case it already used the ISC method. Otherwise, we just referred to the more recent work adopting ISC analysis (for an example of that see the ref. Nastase et al., 2020 in Supplementary Table 3). The calculation of the total data points was performed dividing the stimulus length in seconds for the TR (or, when available we just considered the number of acquired functional scans) and multiplying the results by the sample size. Note that, since our aim was to recruit a sufficient number of sensory deprived subjects to guarantee reliability of ISC results, the total number of data points of the present work was calculated considering the sample size of blind/deaf participants (n=9; therefore, the orange dot indicates log10 of 1614 functional scans by 9 subjects), although the ISC was performed across the two groups (n=18).

### Supplementary Tables

| Cortical Area Label | ROI ID (Glasser et al., 2016) |
|---|---|
| V1 | 1 |
| V2 | 4 |
| V3 | 5,13, 19, 158 |
| V4 | 6 |
| V8 | 7 |
| V7 | 16 |
| V6 | 3, 152 |
| PIT (Posterior InferoTemporal) | 22 |
| FFC (Fusiform Face Complex) | 18 |
| PH (Temporal Cortex) | 138 |
| PHT (Middle Temporal Gyrus) | 137 |
| PHA (ParaHippocampal Cortex) | 126, 127, 155 |
| hMT (human MT) | 2, 23 |
| Early auditory | 24,104, 124, 173, 174 |
| A4 | 175 |
| A5 | 125 |
| STV (Superior Temporal Visual Cortex) | 28 |
| PSL (PeriSylvian Language) | 25 |
| PI (ParaInsular) | 178 |
| STGa (Superior Temporal Cortex - anterior) | 123 |
| STGd (Superior Temporal Cortex - dorsal) | 128, 129 |
| STSv (Superior Temporal Cortex - ventral) | 130, 176 |
| TGd (Lateral Temporal Cortex - dorsal) | 131 |
| TA2 (Auditory Association) | 107 |
| TPOJ (Temporo-Parieto-Occipital Junction) | 139, 140, 141 |
| TE1 (Middle Temporal Gyrus) | 132, 133, 177 |

| | |
|---|---|
| TE2 (Inferior Temporal Sulcus/Gyrus) | 134, 136 |
| Area 7 (Parietal Cortex) | 29, 42, 45, 46, 47 |
| POS (ParietoOccipital Sulcus) | 15, 31 |
| PostCin (Post Cingulate Cortex) | 27, 30, 32, 33, 34, 38, 35, 161, 162, 14 |
| PF (rostral Inferior Parietal Cortex) | 147, 148 149 |
| IPS (IntraParietal Sulcus) | 17,48,49,50, 95,117 |
| IP (IntraParietal Cortex) | 144, 145, 146 |
| PG (intermediate Inferior Parietal Cortex) | 143, 150, 151 |
| IFS (Inferior Frontal Sulcus) | 81, 82 |
| IFJ (Inferior Frontal Cortex) | 79, 80 |
| BA44 (Inferior Frontal, pars Opercularis) | 74 |
| BA45 (Inferior Frontal, pars Triangularis) | 75 |
| Area 47 (Inferior Frontal, pars Orbitalis) | 66, 76, 77, 94, 171 |
| Area 46 (Dorsolateral Prefrontal Cortex) | 83, 84, 85, 86 |
| Area 8 (Medial Prefrontal Cortex) | 63, 67, 68, 70, 73 |
| Area 9 (Medial Prefrontal Cortex) | 69, 71, 87 |
| Area 6-8 (Dorsolateral Prefrontal Cortex) | 97, 98 |
| AntCin (Anterior Cingulate Cortex) | 57, 58, 59, 60, 61, 62, 64, 165, 179, 180 |
| Area 10 (OrbitoFrontal Cortex) | 65, 72, 88, 89, 90, 170 |
| PreMot (PreMotor Cortex) | 12, 54, 96, 10, 11, 56, 78 |

**Supplementary Table 1.** Correspondences between cortical labels and the brain parcellation provided in the HCP Atlas (Glasser et al., 2016).

| Subject | Gender | Age | Cause of blindness | Residual Light Perception | Age of Braille reading |
|---|---|---|---|---|---|
| 1 | M | 49 | retinopathy of prematurity | NLP | 6 |
| 2 | M | 42 | retinitis pigmentosa | NLP | 6 |
| 3 | M | 49 | optic nerve atrophy | NLP | 6 |
| 4 | F | 37 | Leber congenital amaurosis | NLP | 6 |
| 5 | M | 32 | retinal detachment | NLP | 6 |
| 6 | F | 41 | bilateral retinoblastoma | NLP | 6 |
| 7 | M | 19 | retinal detachment | NLP | 6 |
| 8 | M | 57 | retinopathy of prematurity | NLP | 6 |
| 9 | F | 69 | optic nerve atrophy | NLP | 6 |

| Subject | Gender | Age | Cause of deafness | First Language | Hearing aid use |
|---|---|---|---|---|---|
| 1 | M | 24 | hereditary | sign | used during childhood |
| 2 | F | 21 | hereditary | sign | used during childhood |
| 3 | M | 24 | hereditary | sign | used during childhood |
| 4 | M | 26 | hereditary | sign | used during childhood |
| 5 | M | 22 | hereditary | sign | used during childhood |
| 6 | F | 18 | hereditary | sign | used during childhood |
| 7 | F | 28 | Sensorineural hearing loss | sign | currently used |
| 8 | F | 32 | hereditary | sign | used during childhood |
| 9 | F | 22 | hereditary | sign | used during childhood |

**Supplementary Table 2.** Characteristics of congenitally blind and congenitally deaf participants. NLP, No Light Perception; M, male; F, female.

| Authors | Year | Stimulus | Condition | N sub s | Length (min) | TR | Datapoints |
|---|---|---|---|---|---|---|---|
| Hasson U., et al. | 2004 | *The Good, the Bad and the Ugly* | audio-visual | 5 | 0:30:00 | 3000 | 600 |
| Golland Y., et al. | 2007 | *The Good, the Bad and the Ugly* | audio-visual | 12 | 0:16:09 | 3000 | 323 |
| Hasson U., et al. | 2008 | *Curb your enthusiasm* | audio-visual | 12 | 0:27:00 | 2000 | 810 |
| Wilson S. M., et al. | 2008 | *Carrotblanca, Hare Do* | only audio | 12 | 0:14:02 | 2000 | 421 |
| Wilson S. M., et al. | 2008 | *Dripalong Daffy, The scarlet Pumpernickel, Box Office Bunny* | audio-visual | 12 | 0:13:14 | 2000 | 397 |
| Kauppi J.P., et al. | 2010 | *Crash* | audio-visual | 15 | 0:36:00 | 3400 | 635 |
| Hasson U., et al. | 2010 | *The Good, the Bad and the Ugly* | audio-visual | 12 | 0:10:00 | 3000 | 200 |
| Lerner Y., et al. | 2011 | *Pieman* | only audio | 15 | 0:07:00 | 1500 | 280 |
| Nummenmaa L., et al. | 2012 | *When Harry Met Sally and The Godfather* | audio-visual | 16 | 0:24:00 | 1737 | 829 |
| Honey C. J., et al. | 2012 | *narrative story* | only audio | 9 | 0:11:21 | 1500 | 454 |
| Salmi J., et al. | 2013 | *The Match Factory Girl* | audio-visual | 13 | 1:07:00 | 2000 | 2010 |

| | | | | | | | |
|---|---|---|---|---|---|---|---|
| Boldt R., et al. | 2013 | *Postia Pappi Jaakobille (audio-drama)* | only audio | 13 | 0:18:51 | 2500 | 452 |
| Abrams D. A., et al. | 2013 | *music pieces* | only audio | 17 | 0:09:35 | 2000 | 288 |
| Regev M., et al. | 2013 | *Pieman* | only audio | 9 | 0:07:00 | 1500 | 280 |
| Hanke M., et al. | 2014 | *Forrest Gump* | only audio | 18 | 2:00:00 | 2000 | 3600 |
| Lahnakoski J. M., et al. | 2014 | *Desperate Housewives* | audio-visual | 20 | 0:20:00 | 2000 | 600 |
| Bernardi G., et al. | 2014 | *On-board camera F1 clips* | audio-visual | 10 | 0:05:25 | 2500 | 130 |
| Vanderwal T., et al. | 2015 | *Inscapes, Ocean's Eleven* | audio-visual | 22 | 0:07:20 | 2500 | 176 |
| Kauttonen J., et al. | 2015 | *At land* | only video | 12 | 0:14:40 | 2000 | 440 |
| Herbec A., et al. | 2015 | *Sleeping Beauty ballet* | audio-visual | 16 | 0:10:40 | 2000 | 320 |
| Schmälzle R., et al. | 2015 | *real-life political speeches* | only-audio | 12 | 0:07:05 | 2500 | 170 |
| Simony E., et al. | 2016 | *Pieman* | only audio | 36 | 0:07:00 | 1500 | 280 |
| Simony E., et al. | 2016 | *Twilight Zone* | audio-visual | 24 | 0:25:00 | 1500 | 1000 |
| Chen J., et al. | 2016 | *Sherlock* | audio-visual | 22 | 0:48:00 | 1500 | 1920 |
| Jääskeläinen I. P., et al. | 2016 | *The Circus, City Lights* | only audio | 18 | 0:31:32 | 2000 | 946 |
| Lu K. H., et al. | 2016 | *The Good, the Bad and the Ugly* | audio-visual | 9 | 0:11:14 | 2000 | 337 |

| | | | | | | | |
|---|---|---|---|---|---|---|---|
| Chen J., et al. | 2016 | *The Lateness of the Hour from Twilight Zone* | audio-visual | 12 | 0:10:00 | 1500 | 400 |
| Ren Y., et al. | 2017 | *The butterfly circus* | audio-visual | 17 | 0:20:00 | 2200 | 545 |
| Chen J., et al. | 2017 | *Sherlock* | audio-visual | 22 | 0:48:00 | 1500 | 1920 |
| Yeshurun Y., et al. | 2017 | *Pretty Mouth and Green my Eyes* | audio-visual | 19 | 0:11:32 | 1500 | 461 |
| Lahnakoski J. M., et al. | 2017 | *Star Wars, Indiana Jones: Raiders of the Lost Ark; James Bond-Golden eye* | audio-visual | 18 | 0:20:42 | 1800 | 690 |
| Nguyen V. T., et al. | 2017 | *The butterfly circus* | audio-visual | 17 | 0:20:00 | 2200 | 545 |
| Bacha-Trams M., et al. | 2017 | *My Sister's Keeper* | audio-visual | 30 | 1:34:56 | 2000 | 2848 |
| Iidaka T., et al. | 2017 | *Mr. Bean* | only video | 15 | 0:08:25 | 2500 | 202 |
| Yeshurun Y., et al. | 2017 | *Story1, Story2* | only audio | 18 | 0:06:44 | 1500 | 269 |
| Schmälzle R., et al. | 2017 | *movie clips* | audio-visual | 24 | 0:04:35 | 2500 | 110 |
| Jang C., et al. | 2017 | *4 video clips* | only video | 15 | 0:12:04 | 2000 | 362 |
| Thomas R.M., et al. | 2018 | *actions routines movie* | only video | 22 | 0:22:00 | 721 | 1831 |
| Haufe S., et al. | 2018 | *Dog Day Afternoon* | audio-visual | 11 | 0:10:50 | 1500 | 433 |
| Pollick F.E., et al. | 2018 | *13 &32* | audio-visual | 18 | 0:05:08 | 2000 | 154 |

| | | | | | | | |
|---|---|---|---|---|---|---|---|
| Parkinson C., et al. | 2018 | *14 clips* | audio-visual | 42 | 0:36:25 | 2000 | 1093 |
| Lankinen K., et al. | 2018 | *At land* | only video | 8 | 0:15:00 | 2000 | 450 |
| Finn E.S., et al. | 2018 | *narrative story* | only audio | 22 | 0:21:50 | 1000 | 1310 |
| Guntupalli J.S., et al. | 2018 | *Raiders of the Lost Ark* | audio-visual | 11 | 2:00:00 | 2500 | 2880 |
| Bacha-Trams M., et al. | 2018 | *My Sister's Keeper* | audio-visual | 26 | 1:34:56 | 2000 | 2848 |
| Nguyen M., et al. | 2019 | *narrative story* | only audio | 18 | 0:07:00 | 1500 | 280 |
| Loiotile R., et al. | 2019 | *Pieman, Taken, Blow Out, The Conjuring* | only audio | 18 | 0:27:02 | 2000 | 811 |
| Blank I. A., et al. | 2019 | *Pieman* | only audio | 19 | 0:07:00 | 2000 | 210 |
| Di X., et al. | 2020 | *Partly Cloudy* | audio-visual | 29 | 0:05:36 | 2000 | 168 |
| Blank I.A., et al. | 2020 | *Pieman* | only audio | 20 | 0:07:00 | 2000 | 210 |
| Visconti di Oleggio Catello M., et al. | 2020 | *The Grand Budapest Hotel* | audio-visual | 25 | 0:50:00 | 1000 | 3000 |
| Fasano M.C., et al. | 2020 | *Piano Sonata K. 98 by Domenico Scarlatti* | audio-visual | 10 | 0:02:31 | 2000 | 76 |
| Sachs M. E., et al. | 2020 | *Discovery of the Camp; Frysta; Race against the sunset (music)* | only audio | 36 | 0:11:08 | 1000 | 668 |
| Salmi J., et al. | 2020 | *Three Wise Men* | audio-visual | 51 | 0:13:37 | 1900 | 430 |
| Hudson M., et al. | 2020 | *The Conjuring 2; Insidious* | audio-visual | 37 | 3:24:58 | 2600 | 4730 |

| | | | | | | | |
|---|---|---|---|---|---|---|---|
| Chang C. H., et al. | 2020 | *Story (part C)* | only audio | 25 | 0:17:15 | 1500 | 690 |
| Nastase S. A., et al. | 2020 | *Slumlord, Reach for the Stars One Small Step at a time* | audio-visual | 16 | 0:29:26 | 1500 | 1177 |
| Nastase S. A., et al. | 2020 | *It is not the fall that gets you* | audio-visual | 18 | 0:09:45 | 1500 | 390 |
| Nastase S. A., et al. | 2020 | *Pie man (PNI)* | only-audio | 39 | 0:06:57 | 1500 | 278 |
| Nastase S. A., et al. | 2020 | *Running from the Bronx (PNI)* | only-audio | 40 | 0:09:21 | 1500 | 374 |
| Nastase S. A., et al. | 2020 | *I knew you were black* | only-audio | 40 | 0:13:21 | 1500 | 534 |
| Nastase S. A., et al. | 2020 | *The man who forgot Ray Bradbury* | only-audio | 40 | 0:13:57 | 1500 | 558 |
| Schmälzle R., et al. | 2020 | *Bang! You are dead* | audio-visual | 494 | 0:08:02 | 2470 | 195 |
| Skaribas E., et al. | 2020 | *Giselle's solo dance in Act II of Giselle* | only video | 10 | 0:01:30 | 2000 | 45 |

**Supplementary Table 3.** Reviewed datasets summarized by: author and year of publication, stimulus used, condition modality, sample size, stimulus duration and number of datapoints acquired (given the fMRI TR).
